# Integration of Neuroimaging and Histopathology in a Squirrel Monkey Model of Spontaneous Cerebral Amyloid Angiopathy and Amyloid-Related Imaging Abnormalities

**DOI:** 10.64898/2026.10.08.757401

**Authors:** Youssef Z Wadghiri, Suleiman Khan, Muhammad T. Soliman, Jakub Szabo, Audrey Murray, Akash G. Patel, Charles Kingsley, Jody Swain, Pramod N. Nehete, Jelle Veraart, Michele M. Mulholland, William D. Hopkins, Thomas Wisniewski, Henrieta Scholtzova

## Abstract

Cerebral amyloid angiopathy (CAA) is a prevalent cerebrovascular pathology in AD and contributes to amyloid-related imaging abnormalities (ARIA), a major complication of current anti-amyloid immunotherapy. Despite its growing clinical importance, critical gaps remain in the development of reliable biomarkers for antemortem diagnosis of CAA. Squirrel monkeys (SQMs) are a valuable non-human primate model with translational relevance due to their universal propensity to develop abundant CAA and spontaneous ARIA in geriatric animals. The present study employed *R*_2_*, the apparent transverse relaxation rate, which is sensitive to susceptibility-related tissue properties, including iron content and blood oxygenation levels, to monitor age-related brain pathology in SQMs with varying CAA severity. Young, early geriatric, and late geriatric SQMs underwent 7T MRI using a dual-purpose *T*_2_*-w 3D multi-gradient-echo sequence for high-resolution anatomy and quantitative *R*_2_* mapping, alongside *T*_2_-w/FLAIR imaging. Whole-brain and gray matter *R*_2_* increased with age. In contrast, white matter *R*_2_* showed an initial increase followed by a late-life reversal, likely reflecting age-related changes in myelin integrity, tissue microstructure, and tissue water content. MRI-histopathology correlation linked *T*_2_*-w MRI hypointensities to CAA-positive iron-laden vessels associated with perivascular inflammation and, in some cases, hemosiderin deposits. Here we confirmed the occurrence of ARIA-H and ARIA-E events in geriatric animals. SQMs provided a valuable opportunity to examine the underlying pathology of spontaneous ARIA-E, revealing distinctive tissue signatures associated with its manifestation. ARIA-E hyperintensities were characterized by reactive gliosis and BBB disruption, whereas more advanced lesions exhibited perivascular lymphocytic infiltration and evidence of myelin and axonal injury. The current study presents compelling evidence linking *R*_2_* alterations with CAA severity and ARIA-related tissue injury, providing a foundation for further advancing early detection of CAA, ARIA risk assessment, and treatment strategies.

## Introduction

Cerebral amyloid angiopathy (CAA), characterized by amyloid-β (Aβ) deposition in the walls of cortical and leptomeningeal vessels, is common in the elderly and present in most Alzheimer’s disease (AD) cases, with greater CAA severity independently associated with cognitive decline [1–6]. Aβ-targeting immunotherapies have reached important milestones, including U.S. Food and Drug Administration approval for early AD, but their clinical benefits remain limited and are often accompanied by CAA-related complications, namely amyloid-related imaging abnormalities (ARIA). ARIA appears as regions of edema or effusion (ARIA-E) in brain parenchyma or sulci, and as microhemorrhages or cortical superficial siderosis (ARIA-H) [7–9]. Although most ARIA events are asymptomatic, some can be severe or fatal [8, 10–12]. Moreover, spontaneous ARIA can occasionally appear as a consequence of underlying CAA in the absence of anti-amyloid therapy [7, 8, 13–15]. Neuroimaging plays an important role in diagnosis and monitoring CAA, yet definitive diagnosis still requires postmortem neuropathological assessment [16–18]. More sensitive biomarkers are therefore urgently needed, not only for earlier detection before irreversible hemorrhagic injury occurs, but also to minimize ARIA risk and improve treatment outcomes. Progress in advancing biomarker discovery has been limited by the lack of animal models that more closely recapitulate sporadic human disease. Most rodent models require genetic or experimental manipulation and do not naturally develop AD/CAA-related pathology, limiting their value for studying spontaneous disease trajectories and for longitudinal biomarker validation [19–21]. Non-human primates (NHPs) offer key advantages, including longer natural lifespans, a human-identical Aβ sequence, abundant white matter (WM), and complex arterial architecture resembling the human brain [20, 22–24]. Squirrel monkeys (SQMs, *Saimiri boliviensis boliviensis*) have emerged as a valuable NHP model for AD and AD-related dementias (ADRD) research due to their age-dependent accumulation of extensive CAA pathology and the occurrence of spontaneous ARIA-E and ARIA-H events in geriatric animals [25–27]. Our prior work established SQMs as a translationally relevant model for evaluating novel therapeutic interventions [28–30]. Hence, this naturally occurring phenotype, together with the close temporal alignment achievable between antemortem *in vivo* imaging and postmortem neuropathology, positions the SQM as a unique platform to study neuroimaging biomarker trajectories associated with CAA severity and ARIA in a large-brained species amenable to clinical-grade MRI.

Brain iron accumulation has been recognized as a feature of aging and neurodegeneration, and has been hypothesized to contribute to oxidative stress and cell death [31–38]. Altered brain iron homeostasis is well documented in AD, with elevated iron and increased transverse relaxation rate (*R*_2_) and its susceptibility-sensitive counterpart, the apparent transverse relaxation rate (*R*_2_*), observed in neocortical and deep gray matter (GM) regions [36, 39, 40]. Unlike *R*_2_, *R*_2_* additionally reflects spin dephasing from local magnetic field inhomogeneities, such as those induced by iron. Iron dysregulation has also been linked to Aβ aggregation, tau pathology, and cognitive decline [32, 34, 35, 41]. CAA itself is associated with increased brain iron and postmortem work has shown that CAA severity correlates with higher *R_2_* values across cortical and deep GM structures [31, 42]. A recent study indicates that the degree of iron elevation modifies the relationship between CAA and cognitive decline, with CAA linked to steeper decline at higher iron levels [31].

Iron-sensitive MRI techniques, including magnetic field correlation imaging [43, 44], field-dependent relaxation rate increase [45], quantitative *R*_2_/*R*_2_* mapping, and quantitative susceptibility mapping, have consistently shown sensitivity to regional brain iron burden [40, 42, 46–50]. Beyond iron deposition, CAA-related vascular stiffening and impaired cerebrovascular reactivity can elevate *R*_2_ and *R*_2_* through increased regional deoxyhemoglobin and localized hypoxia, a blood oxygenation level-dependent (BOLD) contribution that adds to, rather than replaces, iron-related susceptibility effects [51–54]. *R*_2_* is also influenced by myelin content and microstructural organization, including the orientation of myelinated fibers relative to the main magnetic field [55, 56]. Thus, high-resolution three-dimensional (3D) multi-gradient-echo (MGE) acquisitions can resolve spatially heterogenous *R*_2_* patterns within anatomical subregions, but these patterns should not be interpreted as specific to iron deposition [55, 57].

In the present study, we used high-resolution 7 Tesla (7T) MRI, including 3D isotropic quantitative *R*_2_* mapping derived from MGE acquisition and *T*_2_-weighted fluid-attenuated inversion recovery (*T*_2_-w FLAIR) imaging, to characterize age-related and region-specific iron-sensitive changes in young, early geriatric, and late geriatric SQMs, with longitudinal follow-up in a subset of animals. The MGE acquisition served a dual purpose, providing both the high-resolution anatomical detail needed to refine a species-specific SQM brain atlas beyond previously published atlases [58, 59] and the quantitative data needed for unbiased, atlas-based *R_2_*\* regional analysis. We further integrated neuroimaging findings with brain pathology correlates to characterize *R_2_\** changes and distinct ARIA manifestations. To our knowledge, no prior *in vivo* study has directly linked CAA-related pathology to quantitative *R_2_*\* while monitoring ARIA incidence in an NHP model vulnerable to vascular amyloid pathology. Together, these assessments were designed to evaluate quantitative *R*_2_*, including its reversal signature in WM [46, 60–64], alongside conventional edema-sensitive imaging as complementary, non-invasive candidate biomarkers of CAA-related pathology and vascular fragility, with potential clinical utility, while also providing insights into ARIA susceptibility to improve outcomes for AD immunotherapy.

## Material and Methods

### Experimental Groups

The present study included young (6-8 years of age; n=12), early geriatric (16-18 years of age; early CAA pathology; n=9), and late geriatric (21-23 years of age; advanced CAA pathology; n=12) female squirrel monkeys (SQMs, *Saimiri boliviensis boliviensis*). SQMs have an average lifespan of 20 years in captivity, with a maximum life expectancy of up to 30 years [65–67]. SQMs reproduce effectively in a harem scheme, which consists of single-male/multi-female groups, with ratios reaching as high as one male to 10-12 females [68, 69]. Because breeding colonies are therefore female-predominant, only female SQMs were included in this study. A comprehensive list of subjects who underwent MRI is provided in **Supplementary Table ST1**.

### Care and Housing

SQMs were obtained from the Squirrel Monkey Breeding and Research Resource (SMBRR) at the Michale E. Keeling Center for Comparative Medicine and Research (KCCMR) which is part of the University of Texas MD Anderson Cancer Center (Bastrop, TX). Animals were socially housed in groups of 3 to 14 individuals, in either large pens or quad cages, and maintained on a 12-hour light/dark cycle (lights on at 07:00, lights off at 19:00 hours). Pens (1.5 × 2.5 × 4.5 m) had concrete flooring partially covered with plastic dry-deck material and were equipped with polyvinyl chloride perches at varying heights, a hide box, and a variety of enrichment devices. In quad cages (0.65 × 1.5 × 1.5 m), all dividers were kept open to give each group access to the full unit, and an external tunnel connected the upper and lower tiers. Quad cages were likewise equipped with perches and enrichment devices but had mesh floors instead of concrete and dry-deck material. Animals were assigned to groups based on a preliminary assessment of their mobility and social interaction levels with other monkeys. They were relocated to their designated housing before study onset and allowed to acclimate to the new environment. Animals were fed the New World Primate Diet (Teklad New World Diet 8794N) fresh produce, and ad libitum access to water. Prior to imaging, the SQMs were transported, awake, to the Department of Veterinary Medicine and Surgery (Houston, TX) via a temperature-controlled transport vehicle. While in Houston, animals were socially housed in quad cages and maintained on a 12-hour light/dark cycle with enrichment items provided by the KCCMR. The animals received ad lib food and water. Animals were transferred awake in climate controlled departmental vans to the Small Animal Imaging Facility (Houston, TX) for collection of the MRI scans.

### Ethics Statement

All animal experiments were carried out according to the provisions of the Animal Welfare Act, PHS Animal Welfare Policy, and the principles of the NIH Guide for the Care and Use of Laboratory Animals. All procedures were approved by the Institutional Animal Care and Use Committee (IACUC) at the New York University School of Medicine and the University of Texas MD Anderson Cancer Center.

### Experimental Set Up

A custom SQM setup was skillfully designed in-house to enable careful restraint while considering the non-magnetic requirements of the MRI scanner. A half cylindrical polycarbonate bed lined with molded fabric was used to accommodate the body shape and prevent pressure sores. All limbs were fitted with stockinette before each subject was positioned supine in the homemade sled. The whole body was then covered by a blanket topped with a temperature-controlled recirculating water pad to help maintain optimal body temperature (35 – 37°C). To prevent motion during imaging, the body was secured to the sled with fabric tape. The bed was then advanced into the magnet bore so that the head was positioned at the magnet isocenter, where the static magnetic field (B_0_) is most homogenous and the gradient fields are most linear, and centered within the radiofrequency (RF) volume coil to ensure uniform RF (B_1_) field coverage. This configuration enabled adequate space within the 86 mm inner-diameter RF coil for proper head orientation and mechanical ventilation via intubation of the animals [70].

### Anesthesia Preparation and Monitoring During Imaging

All animals were sedated with a combination of ketamine (10 mg/kg) and xylazine (2 mg/kg) administered intramuscularly. Orotracheal intubation was performed after confirming adequate sedation and desensitizing the larynx with 2% lidocaine (2 mg/kg). Anesthesia was maintained with isoflurane (1-3%) throughout positioning and image acquisition. Lactated Ringer’s solution was administered as a constant rate infusion (1.5 ml/hr) during imaging. Physiological parameters including heart rate, core body temperature, respiration rate via a pressure-sensitive transducer wrapped around the chest, end tidal CO_2_, and SpO_2_ were monitored and maintained at normal levels throughout the imaging sessions with Philips *in vivo* monitoring equipment. During recovery, a single dose of meloxicam (0.1–0.2 mg/kg) and a 30 ml bolus of lactated Ringer’s solution were administered subcutaneously as part of the recovery protocol for each subject. Post-procedure, isoflurane administration was discontinued, and the endotracheal tube was removed shortly thereafter, allowing the animals to recover consciousness. Post-anesthetic recovery was closely monitored to prevent complications.

### High-Resolution MRI Acquisition and Instrumentation

All *in vivo* imaging procedures were performed on a 7T Bruker Biospec 70/30 micro-MRI system (Bruker Biopsin, Billerica, MA, USA), corresponding to the proton (^1^H) Larmor frequency of 300 MHz. The installation is equipped with a superconducting, zero helium boil-off horizontal bore magnet with an inner accessible diameter of 300 mm, based on ultra-shield refrigerated magnet technology. The magnet includes an actively shielded gradient coil insert (Bruker BGA-12S-HP; outer diameter (OD) = 198 mm, inner diameter (ID) = 114 mm, 630 mT/m gradient strength, 130 μs rise time), powered by high-performance gradient amplifiers (IECO, Helsinki, Finland) and operating at 300 A/500 V. The system is interfaced with a Bruker Avance III-HD console running Paravision 6.0.1 and TopSpin 3.1. Imaging setup used a Bruker volume transmit/receive circularly polarized RF resonator (OD = 112 mm, ID = 86 mm, length = 100 mm, model 1P-T12053V3) tuned to 300.16 MHz for scanning SQM subjects.

### MRI Pulse Sequences

Subjects underwent one or two MRI examinations; when two sessions were performed, scans were separated by at least two weeks.

#### Anatomical mapping and MRI sensitive to iron

A 3D *T*_2_*-w MGE dataset was acquired to investigate iron-related pathology. An isotropic spatial resolution of 200 μm was chosen to optimize signal-to-noise ratio (SNR) while maintaining sufficient anatomical detail for inter-session and inter-subject realignment. Pulse sequence parameters included: first echo time (TE) = 2.96 ms, echo spacing (ES) = 4 ms, 8 echoes, repetition time (TR) = 41 ms and a flip angle (FA) of ∼13°, a matrix size (Mx) of 384 x 256 x 256, a field of view (FOV) of 76.8 mm x 51.2 mm x 51.2 mm, and a spectral width (SW) of 125 kHz. The resulting bandwidth (BW) per pixel in the frequency-encoding direction was 325.52 Hz. Acquisition time (T_ACQ_) was 45 minutes per dataset with number of averages (NA) = 1, repeated up to three times (number of repetitions, NR = 3), provided the animal remained physiologically stable under anesthesia, for a total experiment time of up to 150 minutes, including 10–15 minutes for setup and calibration.

#### Edema-sensitive MRI acquisition (T_2_-w/FLAIR)

Edema-sensitive imaging was performed during a separate session scheduled at least two weeks following MGE, thereby allowing sufficient time for complete recovery and subject acclimation. This session consisted of two two-dimensional (2D) multi-slice (MS) pulse sequences acquired sequentially: a *T*_2_-w Turbo-Spin Echo (TSE) and a FLAIR sequence. Both sequences shared the following geometric and acquisition parameters: 200 μm in-plane resolution, 800 μm slice thickness, a 2.4 mm slice gap, Mx = 320 x 320, FOV = 64 mm x 64 mm, and number of slices (NS) = 15. Full brain coverage was achieved for both TSE and FLAIR by acquiring four consecutive 800 µm slice-shifted 2D MS scans to fill in the inter-slice gaps.

TSE acquisition details: The *T*_2_-w TSE sequence (also known as Fast Spin Echo or FSE and as Rapid Acquisition with Relaxation Enhancement or RARE) was acquired using the following parameters: Turbo-Factor (TF) = 10, TE = 7 ms, ES = 7 ms, effective TE (TE_eff_) = 35 ms, TR = 3200 ms, FA = 90°/180°, NA = 4, NR = 4, SW = 81.5 kHz, BW = 254.75 Hz/pixel. The TSE acquisition was comprised of four consecutive 7-minute scans, with a total acquisition time of 28 minutes.

#### FLAIR acquisition details

The FLAIR sequence used the TSE backbone but incorporated an inversion pulse to suppress the cerebrospinal fluid (CSF) signal. It used the same TSE parameters with two exceptions: an inversion time (TI) = 1557 ms was added, and the number of averages was increased to NA = 12 (a threefold increase compared to TSE) to compensate for the signal loss induced by the inversion pulse. Consequently, the FLAIR sequence required four consecutive 20-minute scans, totaling 80 minutes of acquisition time.

### Image Post-Processing

#### Echo-Averaged T_2_*-w MRI and Quantitative R_2_* Mapping

Datasets were registered to the VALiDATe29 digital atlas for cross-sectional and longitudinal assessment of age-related brain changes and atlas refinement.

Combined echo-averaged *T*_2_*-w images (hereafter referred to as echo-averaged) were generated from 3D MGE datasets using MRtrix3 [71]. Echo-averaged *T*_2_*-w images were registered to the VALiDATe29 template in two stages: an initial linear (affine) registration using ITK-SNAP v4.0.2 [72], followed by nonlinear (deformable) registration using ANTs 2.0 [73]. Quantitative, voxel-wise 3D *R*_2_* brain maps were generated using FireVoxel (build 368, https://firevoxel.org/) by fitting a mono-exponential decay model of the apparent transverse relaxation time (*T*_2_*) to the MGE data. For each voxel, the apparent transverse relaxation rate (*R*_2_* = 1/*T*_2_*) was estimated by fitting the signal intensity across the eight echo times to the equation: *S*_(TE)_ = *S*_0_ * exp (-TE _✕_ *R*_2_*), where *S*_(TE)_ is the signal intensity at TE and *S*_0_ is the signal intensity TE = 0. Resulting *R*_2_* maps were used for subsequent analysis. Because the *R*_2_* maps and echo-averaged images were derived from the same MGE acquisition, they were inherently co-registered. Each subject’s *R*_2_* map was therefore transformed into atlas space by sequentially applying the affine transformation and the nonlinear deformation field computed during registration, of the echo-averaged images, using linear interpolation in ANTs. Atlas ROI labels were then overlaid on the transformed maps, and mean *R*_2_* was computed within each ROI using MRtrix3, with the exclusions described below.

#### T_2_-w TSE and FLAIR Images

The *T*_2_-w 2D TSE and FLAIR acquired images were each processed in MRtrix3 by interleaving and concatenating the four sets of slice-shifted 2D MS scans to produce a single full-coverage brain dataset.

### Squirrel Monkey MRI Neuro-atlas

Initial anatomical identification was based on an in-house 3D SQM brain atlas comprising 21 ROIs. Cortical and deep GM boundaries were delineated using a thionin/hematoxylin-stained histological SQM brain atlas (University of Wisconsin and Michigan State Comparative Mammalian Brain Collections, https://brainmuseum.org/index.html) and further refined using resources from Dubach & Bowden (2009) [74] and Rohlfing *et al.* (2012) [75]. WM ROIs were hand-drawn based on the VALiDATe29 and CHIASM MRI SQM brain atlases [58, 59].

To preserve the accuracy of quantitative *R*_2_* measurements, inferior and posterior portions of the temporal and occipital lobes were excluded from analysis. These regions are affected by two independent sources of artifacts in *T*_2_*-w imaging: (i) pronounced physiological motion transmitted from the neck and thorax during respiration, and (ii) severe magnetic susceptibility-induced distortion and signal dropout caused by proximity to the air-filled paranasal sinuses and interaural cavities.

Segmentation masks for total GM and total WM were derived by combining their respective atlas-based subregions. The whole-brain mask was then generated as the union of the total GM and total WM masks for subsequent voxel-based quantification.

### Tissue Collection and Histological Analysis

#### Euthanasia and Brain Tissue Processing

SQM euthanasia and brain tissue processing were conducted as previously described [30, 76]. Subjects were anesthetized with sodium pentobarbital (150 mg/kg, intraperitoneally) and subsequently euthanized for analysis of brain pathology. Transcardial perfusion was performed using 0.1 M phosphate-buffered saline (PBS) supplemented with heparin solution (10,000 U/L; pH 7.4). Following perfusion, the brain was removed and the cerebellum was dissected prior to coronal sectioning. The brain was then cut rostrocaudally into 4-mm-thick coronal blocks using a commercially available SQM brain slicer (Zivic Instruments). To separate hemispheres, coronal blocks were cut along the sagittal midline. In the present study, the right hemisphere was immersion-fixed in 10% neutral-buffered formalin for 48 to 72 hours at 4°C. For selected geriatric animals, both hemispheres were fixed when available. Following fixation, brain blocks were processed for paraffin embedding and subsequent histological sectioning (NYU Langone Experimental Pathology Research Laboratory).

MRI-histopathology correlation was restricted to late-geriatric subjects, because animals in the young and early-geriatric cohorts were retained for the colony breeding program. Brain tissue from two additional subjects, age-matched to the young (6-year-old) and early-geriatric (17-year-old) cohorts, was also examined to document age-related differences in amyloid burden. These subjects were not part of the current imaging protocol.

#### Immunohistochemistry and Histology

Formalin-fixed, paraffin-embedded coronal brain sections cut at a thickness of 8 µm were used for histological assessment of brain pathology utilizing standard techniques [25, 26, 30]. Immmunostaining was performed using standard protocols and as below [25, 26, 30, 76–78]. Parenchymal and vascular Aβ deposits (CAA) were detected with a mixture of 6E10 and 4G8 anti-Aβ antibodies (total amyloid). IBA1 (ionized calcium-binding adapter molecule 1) was used for the detection of the entire microglia population (resting, activated states) [79, 80]. Glial fibrillary acidic protein (GFAP) served as a marker of astrocytic activation. CD163 immunohistochemistry was used to identify perivascular macrophages [81]. T lymphocytes were identified by CD3 immunoreactivity. Two markers were used to assess blood-brain barrier (BBB) breakdown. Antibody for fibrinogen, a plasma glycoprotein essential for clot formation, was used to detect parenchymal and intravascular fibrinogen gamma chain (FGG). Immunoglobulin G (IgG) immunohistochemistry was performed to further confirm disruption of the BBB. Immunostaining for myelin basic protein (MBP), a major structural protein of the myelin sheath, was used to assess myelin integrity and detect demyelination. Phosphorylated neurofilament heavy chain (SMI-31) was used as a marker for axonal integrity [26]. Ferritin heavy chain (ferritin) immunostaining was used to label the iron-storage protein ferritin. Antigen retrieval was required for the detection of 6E10/4G8 (88% formic acid), IBA1 (10 mM Tris/1 mM EDTA buffer, pH 9.0), CD163 (10 mM Tris/1 mM EDTA buffer, pH 9.0), fibrinogen (10 mM sodium citrate buffer, pH 6.0), IgG (proteinase K [PK] 1:100 - 10 mM Tris/1 mM EDTA buffer, pH 8.0), CD3 (10 mM sodium citrate buffer, pH 6.0), MBP (10 mM sodium citrate buffer, pH 6.0), ferritin (PK 1:100 – 10 mM Tris/1mM EDTA buffer, pH 8.0), and SMI-31 (10 mM sodium citrate buffer, pH 6.0) immunostainings. Endogenous peroxidases were blocked by incubating in 0.3% H_2_O_2_/PBS (pH 7.4). Sections were then blocked for 2 hours in either horse blocking solution (10% normal horse serum with 0.3% Triton X-100 in PBS) or goat blocking solution (10% normal goat serum with 0.3% Triton X-100 in PBS). Primary anti-Aβ antibodies 6E10 and 4G8 (1:1000), CD3 (1:20), MBP (1:1000), SMI-31 (1:1000), IBA1 (1:500), GFAP (1:500), CD163 (1:100), fibrinogen (1:500), IgG (1:200), and ferritin (1:1000), were applied in a primary antibody diluent composed of 0.3% Triton X-100, 0.1% sodium azide, and 3% normal horse serum/normal goat serum in PBS, at 4°C overnight. Primary antibody incubation was followed by incubation with either secondary biotinylated anti-mouse IgG reagent or secondary biotinylated anti-rabbit IgG reagent at room temperature (RT) for 1 hour. Next, sections were incubated with ABC solution for 1 hour at RT. For IgG staining, the primary antibody was directly conjugated to horseradish peroxidase, omitting the secondary antibody step. Antibody staining was revealed with 3,3′-diaminobenzidine (DAB) with nickel ammonium sulfate intensification. A hematoxylin counterstain was applied following the completion of fibrinogen and ferritin immunohistochemical staining. Nuclear Fast Red (Ricca Chemical Co., Arlington, TX) was applied following the completion of IgG and CD3 immunohistochemical staining.

Hemosiderin associated with microbleeds was detected using Perls’ Prussian blue staining, and was performed as previously described [30, 76, 77]. Briefly, sections were incubated in an 8% potassium hexacyanoferrate and 8% hydrochloric acid solution in deionized water (DI-H_2_O) at 37°C for 30 minutes. Nuclear Fast Red counterstaining was then applied. Hematoxylin & eosin (H&E) and Luxol Fast Blue and eosin (LFB) staining were performed according to standard histology protocols [26].

Double staining for ferric iron (Fe^3+^) and either Aβ deposits (6E10/4G8) or microglia (Iba1) was performed using the Meguro iron staining method [82, 83]. Briefly, sections were incubated in a 1% solution of potassium hexacyanoferrate adjusted to pH 1.3 at 60°C for 30 minutes. Incubation was followed by treatment with a 0.01 M NaN_3_ and 0.03% H_2_O_2_ methanol solution for 1 hour. Iron staining was intensified with DAB in PBS for up to 30 minutes. Following intensification and antigen retrieval, sections were blocked and incubated overnight at 4°C with either 6E10/4G8 or IBA1 primary antibodies, followed by the appropriate biotinylated secondary antibody reagent at RT for 1 hour. Visualization was achieved using VECTASTAIN® Elite ABC-AP Kit and Vector® Red Alkaline Phosphatase Substrate Kit according to manufacturer’s protocol. All stained coronal sections were scanned using an Aperio VERSA 8 (Leica Biosystems, Deer Park, IL) microscope slide scanner for subsequent histopathological analysis.

### Statistical Analysis

Statistical analyses were performed with IBM SPSS Statistics 28.0 (IBM, Armonk, NY) and data were plotted using GraphPad Prism 10.4.0. (GraphPad Software, San Diego, CA). An a *priori* power power analysis (G*Power 3.1.9.7, one-way ANOVA, across three age cohorts, α = 0.05, power = 0.80, large effect size f = 0.6) indicated a required sample size of 30 animals. Because the number of animals available from the breeding colony was limited, the number of animals with *R*_2_* data (n=23) fell below this target. However, replicate MRI acquisitions per subject improved the precision of individual *R*_2_* estimates but did not increase the number of independent subjects. *T*_2_*-w MGE data were acquired with up to three repetitions per animal; fewer repetitions were available for some animals, and each animal’s *R*_2_* value was therefore weighted by the number of repetitions acquired. Normality was assessed by consensus of the Shapiro-Wilk, D’Agostino-Pearson and Anderson-Darling tests. *R*_2_* values of the three age cohorts were compared cross-sectionally using One-way analysis of variance (ANOVA), followed by Bonferroni-corrected post-hoc comparisons when the ANOVA was statistically significant. A subset of three SQMs was followed longitudinally, and *R*_2_* values were compared within-subjects using two-tailed paired *t*-tests (or Wilcoxon signed-rank tests when normality was not met). Moreover, Pearson’s correlation coefficient (r) was calculated to assess the relationship between age and *R*_2_* values for all SQMs in the whole brain and total GM.

Because WM *R*_2_* followed a non-linear age trajectory, linear and quadratic regression models were compared using the Akaike information criterion (AIC) and adjusted R² to select the better-fitting model. Data are reported as mean ± standard error of the mean (SEM) with 95% confidence intervals, and *p* ≤ 0.05 was considered significant.

## Results

The MGE acquisition served a dual purpose, providing both high-quality anatomical guidance and quantitative *R*_2_* mapping in a single scan. SNR declined across the eight echoes as expected from *T*_2_* decay, with early echoes yielding higher SNR but lower GM/WM contrast, and later echoes providing greater contrast at the cost of SNR (**Supplementary Figure S2A**). Multi-echo averaging exploited this complementarity, boosting GM SNR by ∼184% relative to the first echo alone and increasing GM/WM CNR by 279–380% relative to echoes 1 and 8, respectively, while also mitigating *T*_2_*-related signal loss near air/tissue interfaces such as the interaural cavities. The resulting multi-echo-averaged volumes afforded superior tissue contrast (**Supplementary Figure S2B**) that facilitated fine anatomical delineation and the definition of granular sub-anatomical regions that aligned with and extended areas in the published atlas [58], culminating in a refined SQM atlas (**Supplementary Figure S2D**). In parallel, the multi-echo train enabled pixel-wise *R*_2_* mapping (**Supplementary Figure S2C**), yielding a quantitative iron-sensitive biomarker that, unlike qualitative contrast-dependent measures, supports unbiased cross-cohort comparisons alongside the high-resolution anatomical reference.

### Cross-sectional *R*_2_* Evaluation of Atlas-Based and Age-Related Differences in SQMs

Whole brain *R*_2_* analysis, encompassing all subjects including the three longitudinally monitored individuals (identified by color-coded paired symbols) is presented in **Supplementary Figure S3**. Pearson’s correlation coefficient revealed a significant, age-related increase in whole brain *R*_2_* (*r* = 0.390, *p* = 0.049, *R^2^* = 0.152; **Supplementary Figure S3A**). This age-related increase was consistently observed in the longitudinal data from the three individually monitored subjects, suggesting that the effect is not driven by inter-subject variability. Whole GM *R*_2_* exhibited a similar, statistically significant, age-related increase (*r* = 0.448, *p* = 0.022, *R^2^* = 0.201; **Supplementary Figure S3B**), WM *R*_2_* showed no linear association with age (r = −0.02, *p* = .88) and followed a curvilinear trajectory instead, with an initial increase followed by *R*_2_* reversal at later ages. A quadratic model fit the data better (ΔAIC ≈ 4.9; adjusted R² = 0.19 vs. −0.05) and was significant overall [F(2, 20) = 3.50, *p* = 0.049, R² = 0.26] and adding the quadratic term significantly improved model fit, [F(1, 20) = 6.98, *p* = .016]. Group-wise cross-sectional comparisons in the subset of animals with *R*_2_* data (n=23 of 33), across young (n=5 of 12), early geriatric (n=6 of 9), and late geriatric (n=12 of 12) cohorts confirmed these age-related trends for whole-brain, total GM, and total WM *R*_2_* (all *p <* 0.001; **Supplementary Figure S3C-E;** full statistics in **Supplementary Table ST3**). In post-hoc comparisons, whole-brain *R*_2_* was significantly higher in late-geriatric than young cohorts (\*\*\**p <* 0.001; **Supplementary Figure S3C**), also exhibiting a significant increase between young and early-geriatric groups (\*\*\**p* = 0.001; **Supplementary Figure S3C**). Total GM *R*_2_* followed the same pattern (\*\*\**p* < 0.001, \**p =* 0.036 respectively; **Supplementary Figure S3D**), while total WM *R*_2_* was significantly elevated in early-geriatric relative to young animals (\*\**p =* 0.002; **Supplementary Figure S3E**) before declining toward late-geriatric age (\*\*\**p* < 0.001; **Supplementary Figure S3E**), consistent with the quadratic trajectory in panel B. Longitudinal trajectories from the three individually monitored SQMs corroborated the cross-sectional patterns, showing significant progressive increases in the whole-brain and GM *R*_2_* together (\*\**p* = 0.002*, ***p* < 0.001 respectively; **Supplementary Figure S3F, G)** with a non-significant late-life reversal in WM *R*_2_* consistent with the quadratic fit described above (**Supplementary Figure S3H**). A complete extent of the longitudinal 3.5-year follow-up quantitative *R*_2_* analysis of the whole brain, total GM, and total WM regions is provided in **Supplementary Table ST4**.

### T_2_*-w and R_2_* Maps

*In vivo T*_2_*-w MRI depicted differences across the three SQM age cohorts, revealing hypointense regions throughout the GM of the whole brain. These regions were visible in the coronal plane along the rostro-caudal axis (**Figure 1**). Examples from the prefrontal (PFC), frontal (FC), parietal-temporal (P-TC), and occipital (OC) cortices are shown in **Figure 1A, 1C, 1E and 1G**, respectively. The presence of increased hypointensities observed with aging corresponded to shorter *T*_2_*, consistent with, but not specific to, elevated iron content. Notably, MRI hypointense regions were categorized into two distinct patterns: 1) continuous hypointense signals following cerebral vessels, likely attributable to inadequate blood flow and insufficient hemoglobin oxygenation, resulting in local hypoxia and a decreased BOLD signal; 2) randomly distributed spots or clusters, likely reflecting microhemorrhages. Corresponding *R*_2_* maps (1/*T*_2_*), displayed in color-coded blue and yellow (**Figure 1B, 1D, 1F, 1H**), highlighted increased *R*_2_* areas (yellow) in geriatric SQM subjects, consistent with iron accumulation within the GM (blue). Note that these maps were generated using arbitrary *R*_2_* thresholds to visually distinguish iron-rich regions (yellow) from surrounding GM (blue). This thresholding enabled clear delineation of the WM regions (yellow) in the young cohort (**Figure 1B, 1D, 1F, 1H**).

**Figure 1:**
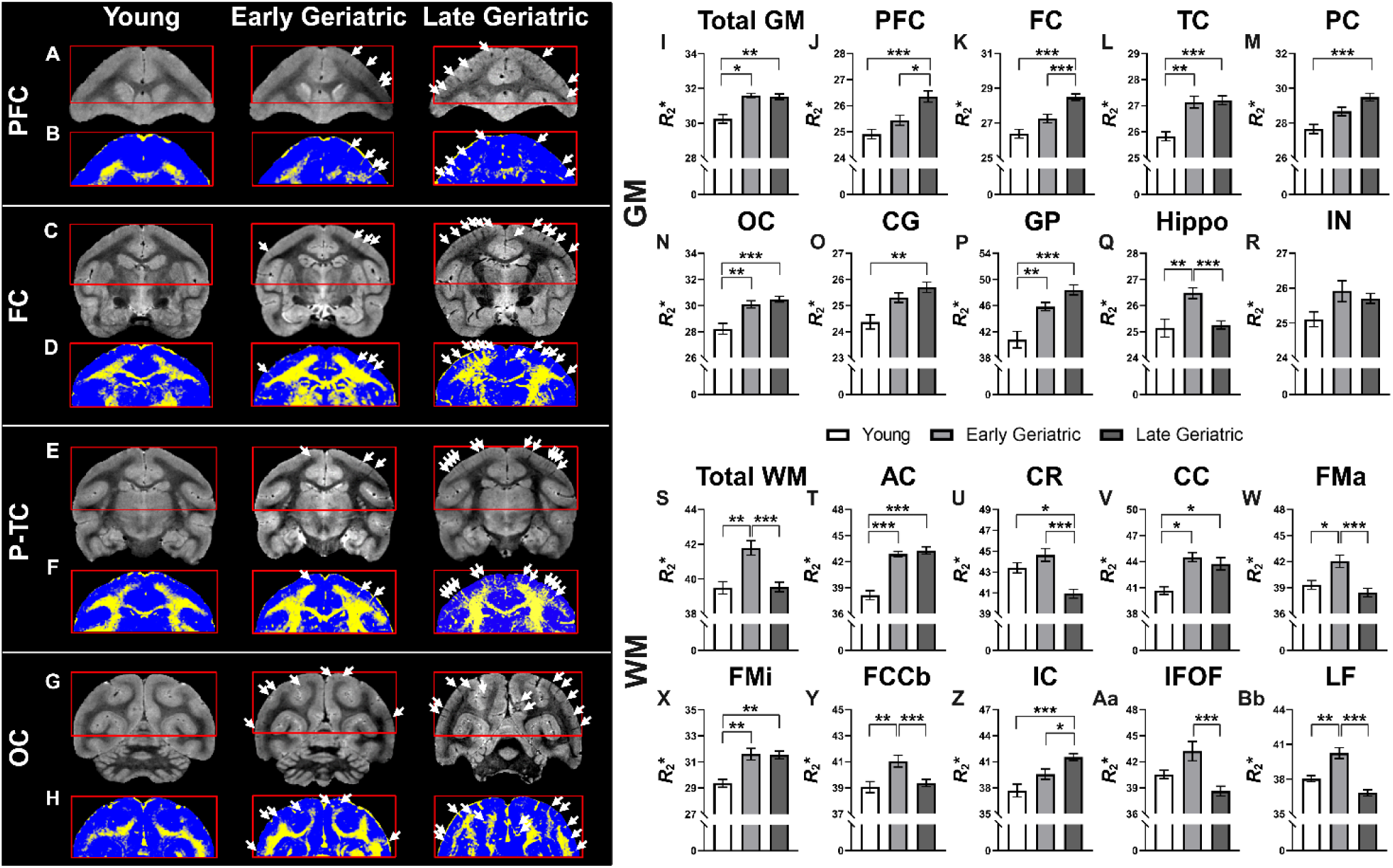
Age-related *R*2* changes across gray and white matter. Left image panel. **(A-H):** Echo averaged *T*_2_*-w images (top row) and corresponding *R*_2_* maps (bottom row) from prefrontal **(A, B)**, frontal **(C, D)**, parieto-temporal **(E, F)**, and occipital **(G, H)** cortices across SQM age cohorts. White arrows indicate progressive GM hypointensities consistent with vessel-associated deoxygenation and microhemorrhages. *R*_2_* maps (blue, GM; yellow, elevated *R*_2_* in WM) confirm age-dependent iron accumulation. Thresholds are applied for visualization only. **Upper right bar-plots panel (GM, I-R):** Four GM *R*_2_* trajectories were observed: continuous increase [**(M)** parietal, **(O)** cingulate]; early-onset increase [**(I)** total GM, **(L)** temporal, **(N)** occipital, **(P)** globus pallidus]; late-onset increase [**(J)** prefrontal, **(K)** frontal]; and increase-then-reversal [**(Q)** hippocampus]. **(R)** No significant age-related changes were detected in insular cortex. **Lower right bar-plots panel (WM, S-Bb):** Three WM *R*_2_* trajectories were observed: early-onset increase [**(T)** anterior commissure, **(V)** corpus callosum, **(X)** forceps minor]; late-onset increase [**(Z)** internal capsule]; and increase-then-reversal [**(S)** total WM, **(U)** corona radiata, **(W)** forceps major, **(Y)** fiber extension of corpus callosum body, **(Aa)** inferior fronto-occipital fasciculus, and **(Bb)** longitudinal fasciculus. Statistical Analysis: One-way ANOVA with Bonferroni correction; see full cross-sectional statistics in **Supplementary Table ST3**. Significance: \**p* ≤ 0.05; \*\**p* ≤ 0.01; \*\*\**p* ≤ 0.001.

### Atlas-based Quantification of R_2_*

Atlas-based *R*_2_* analysis was performed in anatomically defined regions delineated on the echo-averaged images. We identified four distinct profiles of GM *R*_2_* changes associated with aging. The first profile, a continuous increase, was characterized by a significant *R*_2_* increase in the late-geriatric cohort compared to the young cohort. This profile was exhibited in the parietal cortex (PC, \*\*\**p* < 0.001; **Figure 1,M**) and cingulate cortex (CG, \*\**p* = 0.002; **Figure 1O**). The second profile showed an early onset of *R*_2_* increase with age, characterized by a significant *R*_2_* increase in early-geriatric monkeys compared to young monkeys, while no further increase was observed between the early- and late geriatric groups. This profile was observed in the temporal cortex (TC, \*\**p* = 0.002; **Figure 1L**) and occipital cortex (OC, *\*\*p* = 0.002; **Figure 1N**). The third profile was defined by a late onset, exhibiting no significant differences between young and early-geriatric animals, but significantly higher *R*_2_* values in late-geriatric subjects when compared to the early-geriatric cohort. This was demonstrated in the prefrontal cortex (PFC, \*\**p* = 0.025; **Figure 1J**) and frontal cortex (FC, \*\*\**p* < 0.001; **Figure 1K**).

Notably, the hippocampus (Hippo, **Figure 1Q**), exhibited a unique fourth profile: a significant increase in *R*_2_* values was detected when comparing animals in the young cohort to early-geriatric cohort (\*\**p* = 0.002) followed by a significant decrease in late-geriatric monkeys compared to early-geriatric animals (\*\*\**p* < 0.001). No significant differences were detected between groups in the insular cortex (IN, **Figure 1R**).

In WM, the continuous-increase profile was not observed in any of the subregions examined. Instead, early onset *R*_2_* increases were found in the anterior commissure (AC, \*\*\**p* < 0.001; **Figure 1T**), corpus callosum (CC, \**p* = 0.024; **Figure 1V**), and forceps minor (FMi, \*\**p* = 0.004; **Figure 1X**). Late onset *R*_2_* increases were observed in the internal capsule (IC, \**p* = 0.026; **Figure 1Z**). The fourth profile, characterized by an initial increase followed by reversal, was prevalent in several WM regions: the corona radiata (CR, increase: *p* = 0.590, ns; decrease: \*\*\**p* < 0.001; **Figure 1U**), forceps major (FMa, increase: \**p* = 0.025; decrease: \*\*\**p* < 0.001; **Figure 1W**), fiber extension of corpus callosum body (FCCb, increase: \*\**p* = 0.004; decrease: \*\*\**p* < 0.001; **Figure 1Y**), inferior fronto-occipital fasciculus (IFOF, increase: *p* = 0.168, ns; decrease: \*\*\**p* < 0.001 **Figure 1Aa**), and longitudinal fasciculus (LF, increase: \*\**p* = 0.003; decrease: \*\*\**p* < 0.001; **Figure 1Bb**).

We further extended our regional analysis to deep GM structures. The globus pallidus (GP, **Figure 1P**) followed the early-onset profile with a significant increase in *R*_2_* values in the early-geriatric compared to young stage (\*\**p* = 0.010), and no further increase comparing early- to late-geriatric animals. Although not significant, the thalamus (THL, **Figure S4A**) revealed a trend similar to the early onset, and the caudate nucleus (CN, **Figure S4B**) exhibited a continuous increase resembling the first profile. On the other hand, the putamen (PU, **Figure S4C**) showed a profile distinct from all other regions, with an initial decline from young to early-geriatric stage followed by an increase from the early- to late-geriatric stage possibly suggesting a third profile. A comprehensive analysis of cross-sectional GM and WM *R*_2_* changes is provided in **Supplementary Table ST3**.

### Longitudinal *R*_2_* Monitoring in Geriatric SQMs Mirrors Cross-Sectional Regional Trajectories

#### T_2_*-w and R_2_* Maps

Three geriatric SQMs underwent longitudinal follow-up MGE scans 3.5 years following their initial scans (17 to 21, 19 to 23, and 19 to 23 years). Longitudinal follow-up multi-echo scans (shown as *T*_2_*-w echo-averaged anatomical images) (**Figures 2A, 2C, 2E, 2G**) and their respective generated *R*_2_* maps (**Figures 2B, 2F, 2F, 2H**) showed more abundant GM hypointensities (white arrows) and corresponding *R*_2_* elevations throughout the brain compared with baseline scans.

**Figure 2:**
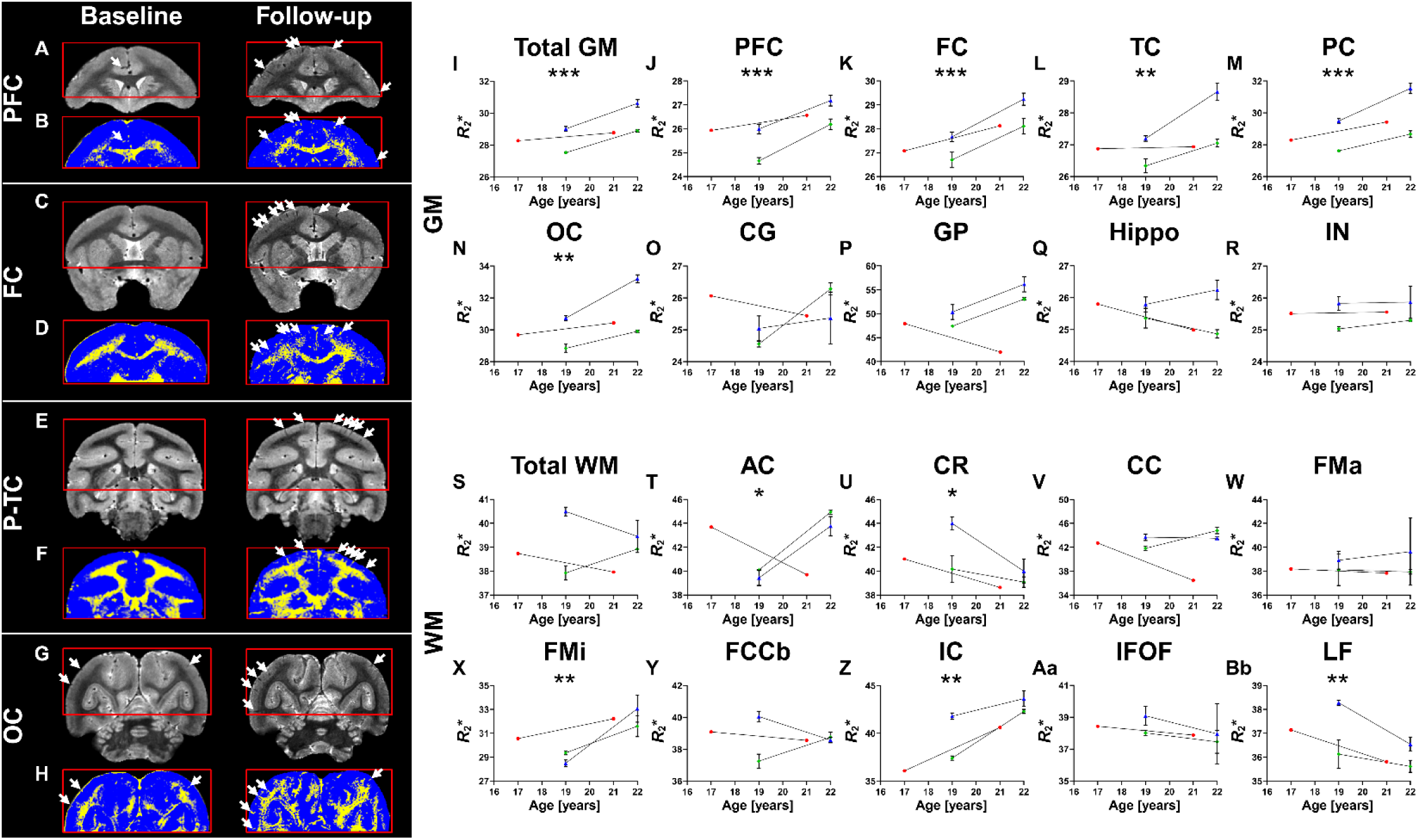
Longitudinal corroboration of age-related *R*_2_* regional trajectories in geriatric SQMs over 3.5 years. **Left image panel** (**A-H):** Echo averaged *T*_2_*-w images (top row) and corresponding *R*_2_* maps (bottom row) from prefrontal **(A, B)**, frontal **(C, D)**, parieto-temporal **(E, F)**, and occipital **(G, H)** cortices across age cohorts. White arrows indicate increased GM hypointensities at follow-up, consistent with vessel-associated deoxygenation and microhemorrhages. Corresponding *R*_2_* maps (blue, GM; yellow, elevated *R*_2_* in WM) confirmed age-dependent iron accumulation at 3.5-year follow-up in three geriatric SQMs (17→21, 19→23, and 19→23 years). Thresholds are applied for visualization only. **Upper right bar-plots panel** (**GM, I-R):** Longitudinal GM *R*_2_* changes: significant *R*_2_* increases were observed in prefrontal, frontal, temporal, parietal, and occipital cortices; no significant *R*_2_* changes were observed in insula, cingulate, and globus pallidus, suggesting a plateau at an early-onset stage. Two of three subjects showed hippocampal *R*_2_* decline, consistent with the reversal profile observed cross-sectionally. **Lower right bar-plots panel** (**WM, S-Bb):** Longitudinal WM *R*_2_* changes: significant *R*_2_* increases were observed in forceps minor, internal capsule, and anterior commissure; significant *R*_2_* declines were detected in corona radiata and longitudinal fasciculus, consistent with the increase-then-reversal trajectory identified cross-sectionally. Statistical Analysis: Paired *t*-test throughout; see full longitudinal statistics in **Supplementary Table ST4.** \**p* ≤ 0.05; \*\**p* ≤ 0.01; \*\*\**p* ≤ 0.001.

#### Atlas-based Quantification of R_2_*

Quantitative *R*_2_* analysis of GM regions over the 3.5-year period corroborated the *R*_2_* increases observed cross-sectionally. Paired *t*-tests demonstrated a significant increases from baseline to follow-up in the prefrontal cortex (PFC, \*\*\**p* < 0.001; **Figure 2J**), frontal cortex (FC, \*\*\**p* < 0.001; **Figure 2K**), temporal cortex (TC, \*\**p* = 0.006; **Figure 2L**), parietal cortex (PC, \*\*\**p* < 0.001; **Figure 2M**), and the occipital cortex (OC, \*\**p* = 0.002; **Figure 2N**). No significant differences were observed in the insula (IN, **Figure 2R**), cingulate cortex (CG, **Figure 2O**), or globus pallidus (GP, **Figure 2P**) between baseline and follow up scans, possibly because *R*_2_* in these regions had already plateaued by the baseline (early-geriatric) scan. The changes among the three subjects in thalamus (THL) were variable (**Figure S4D**) while significantly higher *R*_2_* values were observed in follow up scans compared to baseline scans for the caudate nucleus (CN, \*\**p* = 0.004; **Figure S4E**) and putamen (PU, \**p* = 0.040; **Figure S4F**). In the hippocampus, two of the three subjects showed an *R*_2_* decline between the early-to late-geriatric time points (Hippo, **Figure 2Q**), consistent with the reversal profile. In WM, longitudinal *R*_2_* changes varied by regions: significant increases were observed in the forceps minor (FMi, \*\**p* = 0.005; **Figure 2X**), internal capsule (IC, \*\**p* = 0.004; **Figure 2Z**), and anterior commissure (AC, \**p* = 0.043; **Figure 2T**) between baseline and follow-up timepoints. The longitudinal fasciculus (LF, \*\**p* = 0.010; **Figure 2Bb**) and corona radiata (CR, \**p* = 0.035; **Figure 2U**) showed a significant decline in *R*_2_* between baseline and follow-up scans. A comprehensive analysis of longitudinal GM and WM *R*_2_* changes is provided in **Supplementary Table ST4**.

#### MRI-Neuropathology Correlative Study

A subset of late geriatric SQMs underwent euthanasia within 1 - 3 months following *in vivo* neuroimaging and their brains were subsequently subjected to comprehensive histopathological examination. MRI/histology assessment was restricted to this late geriatric cohort due to the unavailability of young and early geriatric SQM brains. The only young and early-geriatric subjects used for histological analyses were assessed solely for amyloid burden and were not included in our MRI protocol. Sampling of the entire neocortex revealed amyloid deposits detected by 6E10/4G8 immunohistochemistry, predominantly in the form of CAA, characteristic of this species. Vascular Aβ deposits were observed in meningeal and intracortical vessels, with capillaries being particularly vulnerable to CAA (**Figure 3B, C**). Moreover, dyshoric changes, amyloid deposits spreading beyond the vessel wall into the surrounding neuropil, were occasionally detected in microvessels (**Figure 3C**). As expected, histological evaluation depicted a more severe total amyloid burden in late geriatric animals compared to the early geriatric cohort (**Figure 3B, C**). No amyloid pathology was detected in young monkeys (**Figure 3A**). The correspondence between hypointensities detected by high resolution *T*_2_*-w MRI and Aβ histology was evaluated via matched coronal brain sections. In a representative 23-year-old late-geriatric SQM subject, a subset of vessel-like MRI signal hypointensities (**Figure 3D, E**) appeared to correspond to larger CAA-positive vessels visualized by Aβ immunohistochemistry (**Figure 3G, H**), consistent with the impaired blood flow and local hypoxia proposed above. Smaller, presumably non-fibrillar, parenchymal and vascular amyloid deposits (**Figure 3I**), were often less visible on *T*_2_*-w MRI (**Figure 3D, F**).

**Figure 3.**
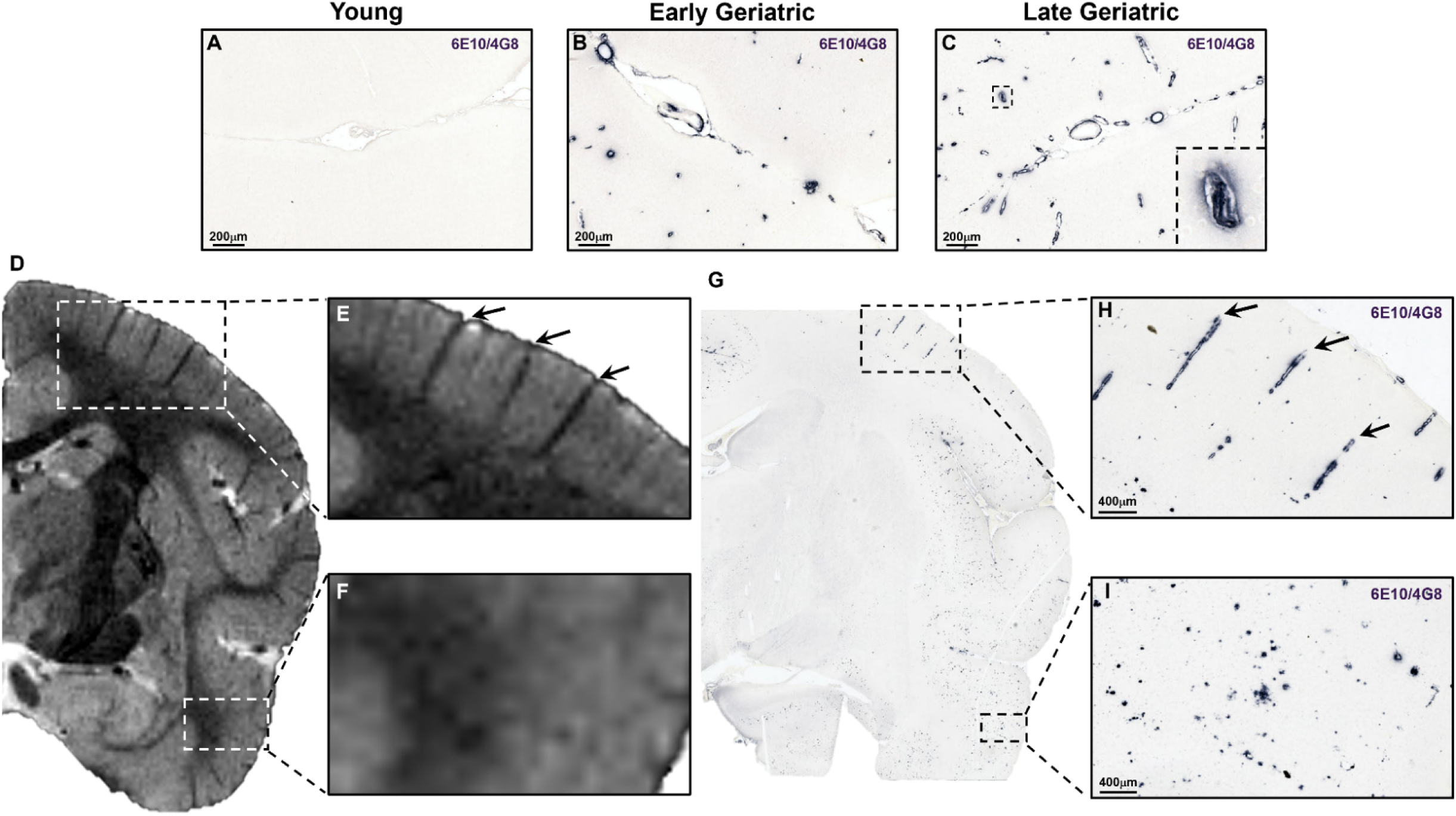
Histopathological examination of CAA and *T*2*-w MRI-histology correspondence in late geriatric SQMs. (A–C) Representative 6E10/4G8 immunohistochemistry across age cohorts: no amyloid in young **(A)**; progressive vascular Aβ burden in early- and late-geriatric animals **(B, C)**, predominantly as CAA in meningeal and intracortical vessels, with capillary involvement and occasional dyshoric changes reflecting amyloid spreading into the surrounding neuropil **(C). (D-I)** MRI-histology correspondence in a representative 23-year-old SQM. Vessel-like *in vivo T*_2_*-w hypointensities **(D, E)** co-localized with larger CAA-positive vessels on Aβ immunohistochemistry **(G, H)**, consistent with hypoperfusion-related susceptibility effects. Smaller, presumably non-fibrillar parenchymal or vascular deposits **(I)** showed limited *T*_2_*-w visibility **(D, F)**, highlighting the resolution-dependent sensitivity of MRI to CAA burden. **(E)** frontal, **(F)** temporal. Scale bars are as indicated.

More comprehensive histological profiling of changes in the vasculature was performed in a 21-year-old geriatric SQM subject. In this example, a CAA-positive penetrating vessel (**Figure 4C**) corresponded to a hypointensity on *T*_2_*-w MRI in the temporal cortex (**Figure 4A, B**). Double staining of an adjacent section using Meguro method for non-heme iron (predominantly present as ferric iron/Fe^3+^) and 6E10/4G8 Aβ immunohistochemistry (CAA) confirmed the iron accumulation within this particular amyloidotic vessel. (**Figure 4D**). This CAA-positive vessel was accompanied by a dense gathering of GFAP immunoreactive astrocytes (**Figure 4E**) and IBA1 immunoreactive microglia (**Figure 4F**). Hence, there appeared to be evidence of perivascular inflammation. Furthermore, fibrinogen immunolabeling revealed the presence of perivascular deposits along with a diffuse pattern in the neuropil (**Figure 4G**), suggestive of BBB disruption. The integrity of this CAA vessel was further assessed using Perls’ Prussian blue staining for ferric reactive hemosiderin deposits; however, no instances of microhemorrhages were detected (**Figure 4H**). Although iron co-accumulation was confirmed within this vessel, the accompanying fibrinogen leakage and dense perivascular gliosis raise the possibility that impaired flow through this amyloid-laden penetrating vessel also contributed a hypoxia-related, BOLD component to the *T*_2_*-w hypointensity; this could not be directly tested in the present example and remains inferential.

**Figure 4.**
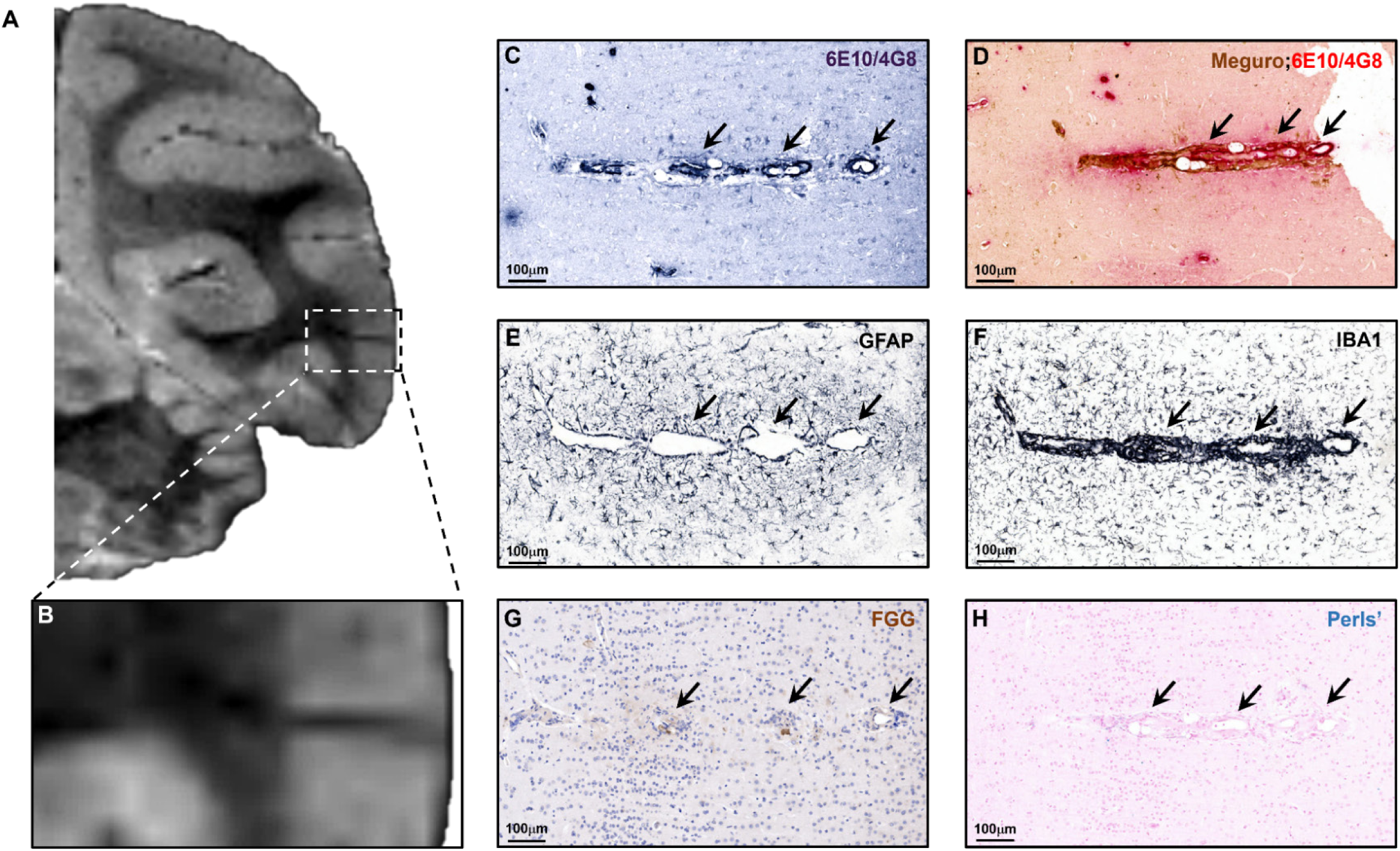
Histopathological characterization of CAA-positive penetrating vessel in a late geriatric SQM. **(A, B)** *In vivo T*_2_*-w hypointense signal enhancement in the temporal cortex corresponded to **(C)** a CAA-positive penetrating vessel confirmed via Aβ immunohistochemistry. **(D)** Double staining of an adjacent section (Meguro iron stain and 6E10/4G8) confirmed iron accumulation within the amyloidotic vessel. **(E, F)** Dense clustering of GFAP-immunoreactive astrocytes and IBA1-immunoreactive microglia, indicating localized perivascular inflammation. **(G)** Fibrinogen immunolabeling revealed perivascular deposits and diffuse neuropil staining, suggestive of increased BBB permeability. **(H)** Perls’ Prussian blue staining showed no evidence of microhemorrhage despite advanced CAA burden. Scale bars, 100 µm.

Several subjects within the late geriatric cohort exhibited ARIA-H signal hypointensities on *T*_2_*-w MRI. One 23-year-old subject with numerous microhemorrhages and cortical superficial siderosis underwent further histological assessment (**Figure 5A**). Hypointense MRI signal alterations in the occipital cortex corresponded to Perls’ Prussian blue-reactive hemosiderin deposits, confirming microhemorrhages and superficial siderosis in the corresponding cortical region (**Figure 5B**). Subsequent double staining using the Meguro method further confirmed iron labeling within vascular amyloid deposits and extravascular iron deposits in the surrounding parenchyma (**Figure 5C**). Additionally, clustering of GFAP-positive astrocytes, IBA1-positive microglia, and CD163-reactive perivascular macrophages was observed in close proximity to ARIA-H affected areas (**Figure 5D-F**). Furthermore, fibrinogen immunohistochemistry demonstrated perivascular deposits, consistent with compromised BBB integrity (**Figure 5G**).

**Figure 5.**
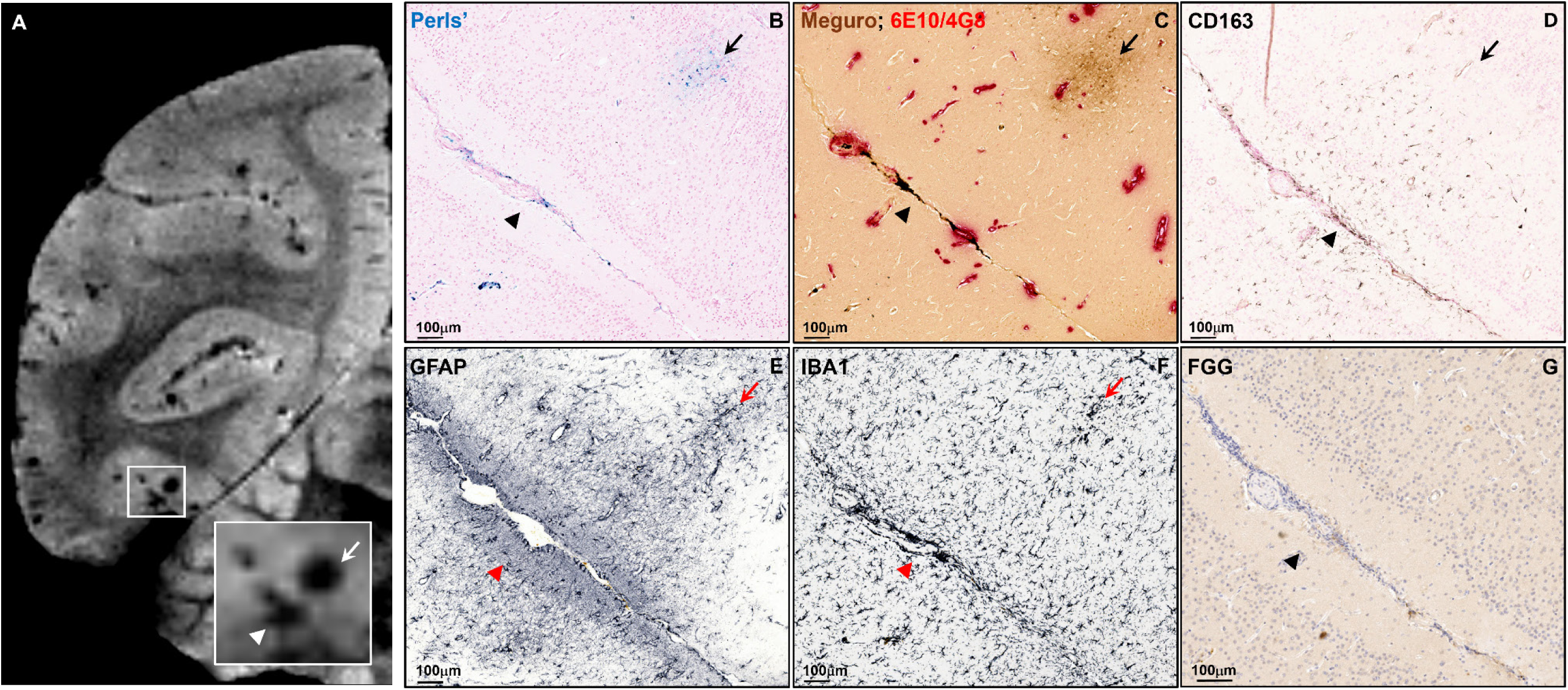
Histopathological characterization of ARIA-H in the occipital cortex of a late-geriatric SQM. **(A)** *In vivo T*_2_*-w MRI showing a focal hypointense signal abnormality in the occipital cortex, consistent with ARIA-H. Inset, higher-magnification view highlighting cortical superficial siderosis and associated microhemorrhagic foci (arrowheads/arrows). **(B)** Perl’s Prussian blue staining of the corresponding cortical region demonstrated hemosiderin deposits, confirming microhemorrhage (arrow) and superficial siderosis (arrowhead). **(C)** Meguro/6E10/4G8 staining identified iron labeling within vascular amyloid deposits and extravascular iron accumulation in the surrounding parenchyma. **(D-F)** GFAP, IBA1, and CD163 immunohistochemistry showed clustering of reactive astrocytes, microglia, and perivascular macrophages respectively, in close proximity to the hemorrhagic area. **(G)** Fibrinogen immunolabeling demonstrated the presence of perivascular deposits. Scale bars, 100 µm.

To further investigate iron-related pathology that may contribute to age-related *R*_2_* changes in SQM brains, we applied two complementary iron-staining approaches. Histological analysis using the Meguro method and IBA1 immunohistochemistry revealed co-localization of microglia with non-heme iron deposits in late-geriatric subjects examined here (**Figure 6B-D**). In one representative 23-year-old geriatric subject, immunohistochemistry for the iron-storage protein ferritin heavy chain identified ferritin-positive cells with morphological features consistent with microglia. These cells were observed in close proximity to a CAA-positive vessel (**Figure 6F, G;** Aβ immunostaining not shown), and IBA1 immunohistochemistry confirmed clustering of microglia around the same vessel (**Figure 6H**). In addition, iron-containing oligodendrocyte-like cells were identified by ferritin immunoreactivity and the Meguro method (**Figure 6B, F**).

**Figure 6.**
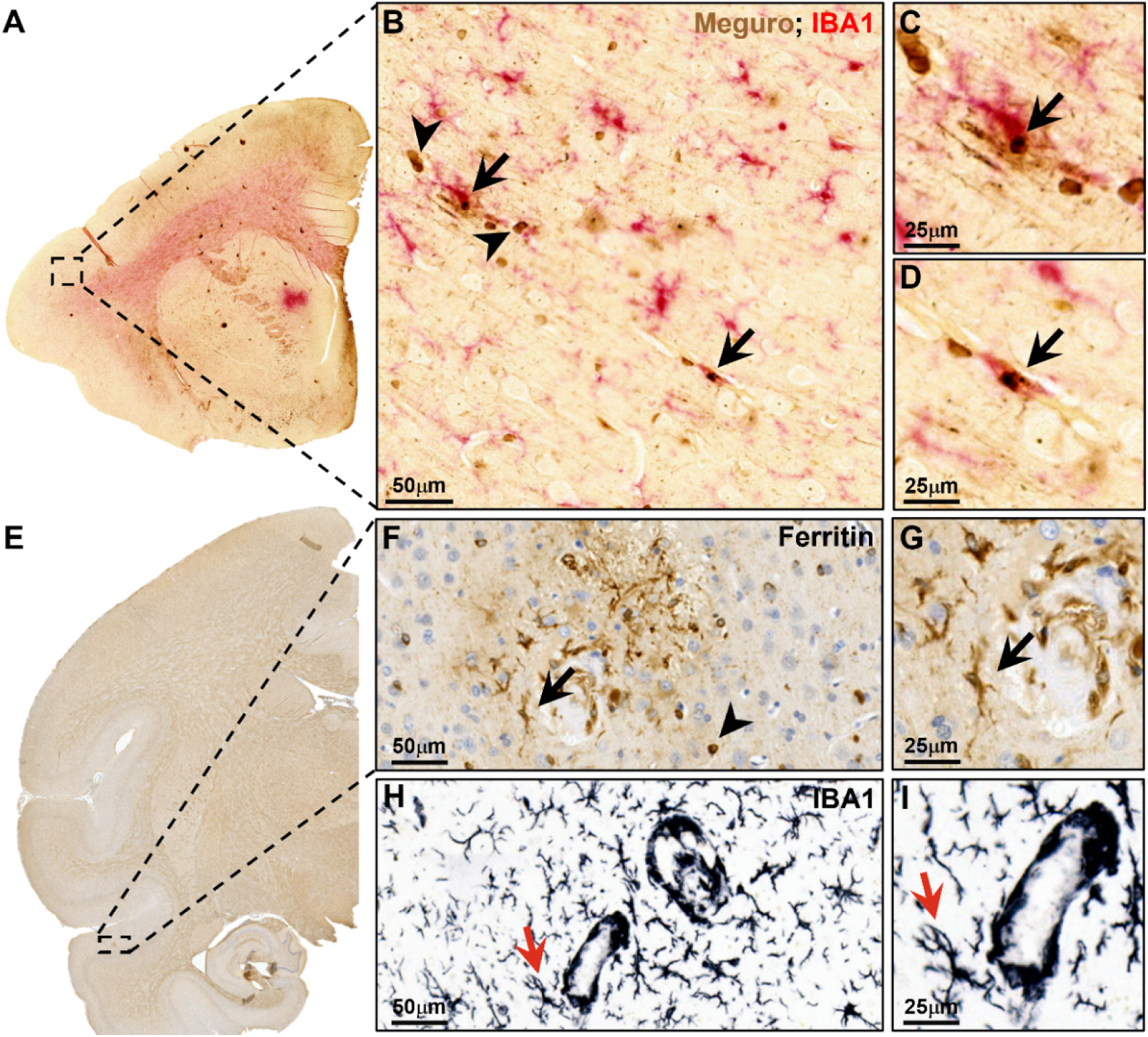
Iron depositions and glial cell-associated pathology in late-geriatric SQMs. **(A)** Low-magnification view of a cortical brain region from a 21-year-old SQM brain processed with the Meguro method and IBA1 immunohistochemistry; the boxed area indicates the region shown at higher magnification in **(B–D)**. **(B–D)** Higher-magnification images demonstrating non-heme iron deposits detected by the Meguro method in close association with IBA1-immunoreactive microglia. Arrows indicate iron-positive deposits co-localized with microglial profiles and arrowheads indicate meguro-positive oligodendrocyte-like cells. **(E)** Low-magnification view of a cortical brain region from a 23-year-old late-geriatric SQM stained for ferritin heavy chain; the boxed area indicates the region shown at higher magnification in **(F–I)**. **(F, G)** Ferritin heavy chain immunohistochemistry identified ferritin-positive cells with morphological features consistent with microglia (arrows) in proximity to a vessel affected by CAA; ferritin-positive oligodendrocyte-like cells are also present (arrowhead). **(H, I)** IBA1 immunohistochemistry of the corresponding region shows clustering of microglia around the same vessel (red arrows). Scale bars are as indicated.

Overall, the quantitative differences in *R*_2_* values observed across our cohorts may reflect a multifactorial substrate involving reduced O₂ availability, impaired perfusion, iron deposition in amyloid aggregates, microhemorrhages, and glial cell/macrophage-associated pathology, several of which were illustrated histologically here.

#### Evaluation of ARIA-E in Squirrel Monkeys

We next screened for hyperintense lesions using a *T*_2_-w 2D TSE and FLAIR imaging. Subcortical hyperintensities, previously attributed to parenchymal vasogenic edema [7], were detected exclusively in late-geriatric SQMs, occurring in 58% of subjects (7/12), whereas no such findings were observed in young (n = 7) or early-geriatric (n = 5) animals. These ARIA-E lesions were identified predominantly in the parieto-occipital lobe, often extending several millimeters along the rostrocaudal axis, with a smaller subset of animals also showing hyperintensities in the frontal and temporal lobes. A summary of ARIA-E MRI findings is provided in **Supplementary Table ST5**.

#### Histopathological Features of ARIA-E Hyperintensities

In a specific 21-year-old subject from the late geriatric cohort, *T*_2_-w TSE MRI revealed the presence of bilateral, multifocal, asymmetric ARIA-E signal abnormalities encompassing the WM (**Figure 7A**) and, to a lesser extent, the GM. The availability of both hemispheres allowed for a more targeted assessment of the histopathological features associated with MRI findings. The first *T*_2_-w hyperintense region was identified in the dorsomedial fronto-parietal WM (**Figure 7A, B**), extending into the adjacent GM (**Supplementary Figure S5A, B**) of the right hemisphere (“***ARIA-E Lesion in Fronto-Parietal Lobe***”). No signal hyperintensities were detected in the corresponding WM of the contralateral hemisphere, which served as the histological control. (**Figure 7A, B**).

**Figure 7.**
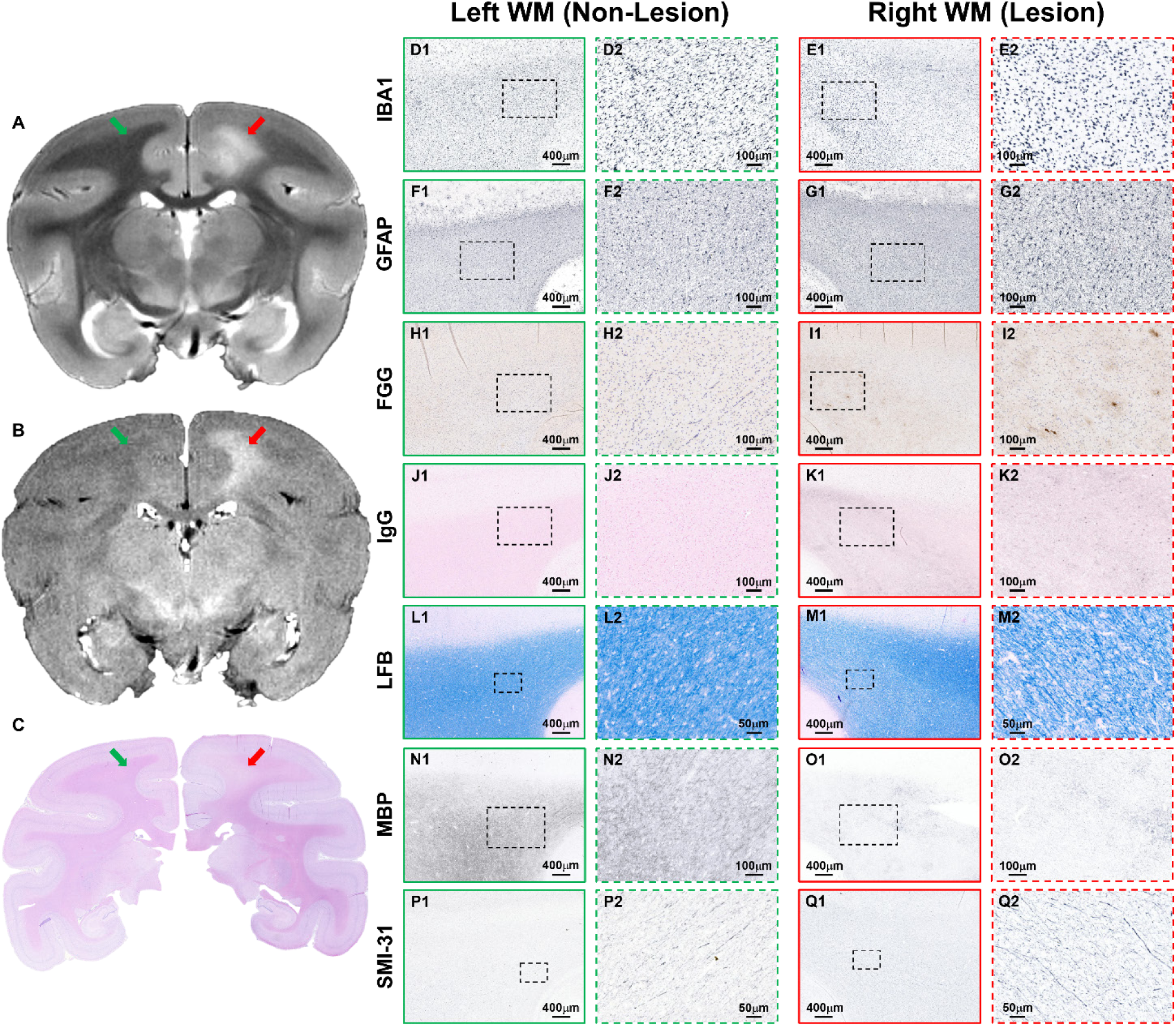
ARIA-E signal abnormality in fronto-parietal WM and histopathological characterization. **(A)** *In vivo T*_2_-w MRI revealed a hyperintense lesion in the dorsomedial fronto-parietal WM of the right hemisphere (red arrow); contralateral normal-appearing WM region (non-lesion WM) is indicated in the left hemisphere (green arrow). **(B)** Corresponding coronal *T*_2_-w FLAIR image confirmed the edematous lesion. **(C)** H&E stain of the matched histological section revealed WM rarefaction. Representative low- and high-magnification sets of histology images are shown for the left hemisphere with normal-appearing WM (non-lesion, green boxes) and contralateral right hemisphere with affected WM (lesion, red boxes), stained for **(D, E)** IBA1, **(F, G)** GFAP, **(H, I)** FGG, **(J, K)** IgG, **(L, M)** LFB, **(N, O)** MBP, and **(P, Q)** SMI-31. No differences in GFAP-positive astrocytes were detected between corresponding lesion and non-lesion WM areas. Relative to contralateral non-lesion WM, the ARIA-E affected WM exhibited dystrophic microglia, enhanced fibrinogen and IgG extravasation, as well as pronounced myelin loss accompanied by axonal abnormalities, consistent with BBB disruption and myelin injury in the setting of ARIA-E hyperintensity. Boxed areas are shown at higher magnification in adjacent panels. Scale bars are as indicated.

H&E staining confirmed the anatomical correspondence with the MRI findings and demonstrated WM rarefaction in the region affected by edema (**Figure 7C**). Histological examination was used to assess the degree of glial activation and focal inflammation in regions with signal abnormalities. There were no apparent differences in IBA1-immunoreactive microglia cells between the lesion (**Figure 7E**) and non-lesion areas (**Figure 7D**). However, IBA1 microglia exhibited dystrophic morphology, with swollen, beaded, and fragmented processes throughout the affected WM region (**Figure 7E**). Similarly, the amount and distribution of GFAP reactive astrocytes appeared comparable between affected (**Figure 7G**) and normal appearing WM (**Figure 7F**). Furthermore, fibrinogen immunolabeling revealed perivascular deposits, along with a diffuse pattern in the ARIA-E-defined WM parenchyma (**Figure 7I**), indicative of BBB disruption. The evidence of plasma extravasation was further supported by multifocal IgG-positive diffuse deposits depicted in this WM lesion area (**Figure 7K**). The MRI-defined ARIA-E-like hyperintense WM region exhibited severe myelin loss, as detected by LFB (**Figure 7M**) and MBP staining (**Figure 7O**), whereas the contralateral WM region lacking apparent signal abnormalities showed only a mild reduction in myelin density (**Figure 7L, N** respectively). The pronounced myelin pallor seen in this region suggests an advanced pathological state. SMI-31 immunohistochemistry for phosphorylated neurofilaments revealed the presence of axonal abnormalities accompanied by sparse spheroids throughout the affected WM area (**Figure 7Q**).

This fronto-parietal ARIA-E lesion e*x*tended into the cingulate sulcus and adjacent GM within the fronto-parietal lobe. A cell-rich area surrounding CAA-positive (6E10/4G8; **Supplementary Figure S5C**) leptomeningeal vessels in the affected sulcus was identified by H&E staining (**Supplementary Figure S5E**). In contrast to the WM findings described above, a prominent astrocytosis (**Supplementary Figure S5G**) and microgliosis (**Supplementary Figure S5I**) were detected in close vicinity to sulcal leptomeningeal vessels, indicative of CAA-associated perivascular inflammation. Furthermore, amyloid-laden small vessels in the surrounding parenchyma were occasionally associated with clusters of reactive astrocytes and microglia (**Supplementary Figure S5G, I** respectively). Additionally, infiltration of CD3-positive lymphocytes was evident within the parenchyma of the edematous GM lesion (**Supplementary Figure S5K**), whereas no CD3-positive T cells were detected in the parenchyma of the contralateral hemisphere (**Supplementary Figure S5J**). Multifocal fibrinogen deposits along microvessels were observed in the affected GM region, consistent with compromised BBB integrity (**Supplementary Figure S5M**).

In the same subject, *T*_2_-w MRI revealed a second ARIA-E like signal abnormality in the dorsolateral occipital WM (**Figure 8A, B**) and GM (**Supplementary Figure S6A, B**) of the left hemisphere (“***ARIA-E Lesion in Occipital Lobe***”). A smaller hyperintense area was identified in the contralateral GM (**Supplementary Figure S6A, B**). IBA1 immunohistochemistry revealed increased microgliosis within the hyperintense WM region of the left hemisphere (**Figure 8D**) compared to the contralateral unaffected WM (**Figure 8E**). No apparent differences in the density of GFAP immunoreactive astrocytes were detected between the lesion and contralateral non-lesion WM areas (**Figure 8F, G** respectively). Extensive fibrinogen extravasation, indicative of BBB dysfunction, was identified in this edematous WM region (**Figure 8H**). IgG halos found around WM microvessels further confirmed the disruption of BBB integrity (**Figure 8J**). No instance of demyelination was detected by LFB or MBP staining in this case (**Figure 8L-O**). Furthermore, a comparable distribution of SMI-31 immunoreactive phosphorylated neurofilament was observed between the lesion and contralateral unaffected WM regions (**Figure 8P, Q**).

**Figure 8.**
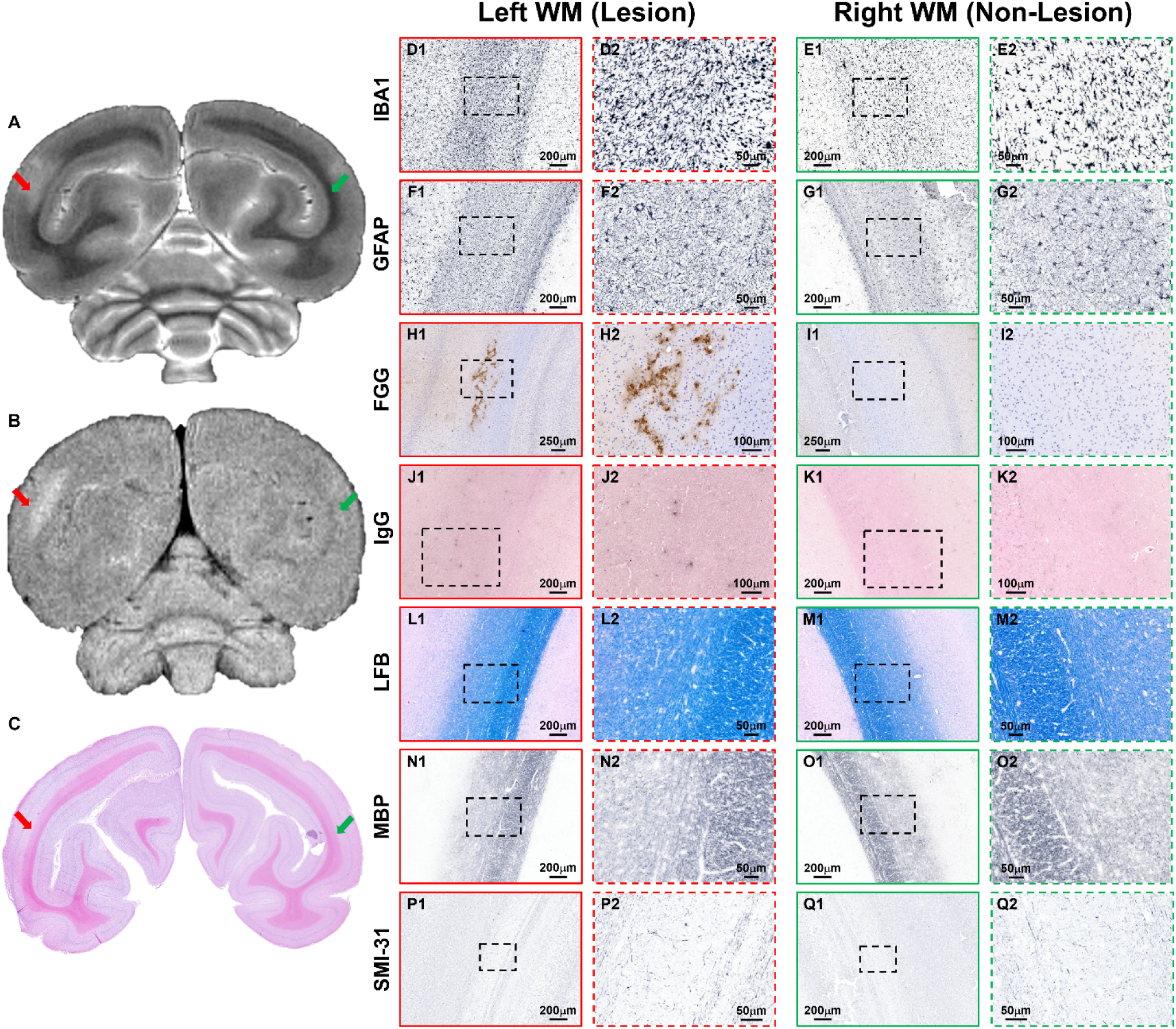
ARIA-E signal abnormality in occipital WM and histopathological characterization. **(A)** *In vivo T*_2_-w MRI showed a hyperintense lesion in the dorsolateral occipital WM of the left hemisphere (red arrow); contralateral normal-appearing WM region is indicated in the right hemisphere (green arrow). **(B)** Corresponding coronal *T*_2_-w FLAIR image confirmed the extent of the edema. **(C)** Matched histological section stained with H&E. Representative low- and high-magnification sets of histology images are shown for left hemisphere with affected WM (lesion, red boxes) and contralateral right hemisphere with normal-appearing WM region (non-lesion, green boxes) stained for **(H, I)** IBA1, (**J, K)** GFAP, **(D, E)** FGG, (**F, G)** IgG, **(L, M)** LFB, **(N, O)** MBP, and **(P, Q)** SMI-31. The left hemisphere affected WM exhibited increased IBA1 immunoreactivity, consistent with reactive microgliosis, whereas GFAP immunoreactivity was comparable between lesion and contralateral non-lesion regions. Marked fibrinogen extravasation and perivascular IgG halos were present in the ARIA-E affected WM, indicating BBB disruption. In contrast, LFB and MBP staining showed no evidence of overt demyelination and SMI-31 immunoreactivity appeared similar between the ARIA-E lesion and contralateral non-affected WM. Boxed areas are shown at higher magnification in adjacent panels. Scale bars are as indicated

We further assessed the histopathological features corresponding to the *T*_2_-w signal hyperintensity in the adjacent occipital GM of the left hemisphere. The hyperintense GM area exhibited increased parenchymal IBA1-positive microgliosis, along with microglia associated with CAA-affected vascular profiles (**Supplementary Figure S6D**), compared to a normal-appearing region in the contralateral hemisphere (**Supplementary Figure S6F**). Pronounced GFAP-reactive astrocytosis, accompanied by perivascular clustering of astrocytes, was also observed in the affected GM region (**Supplementary Figure S6G**) compared to the contralateral unaffected GM (**Supplementary Figure S6I**). Moreover, perivascular fibrinogen-immunoreactive deposits were identified throughout the ARIA-E GM lesion area (**Supplementary Figure S6J**). A second smaller GM lesion, detected in the contralateral right hemisphere, displayed similar histopathological features when compared to unaffected GM regions (**Supplementary Figure S6E, H, K**).

## Discussion

CAA is a common comorbidity in AD and has been implicated in cerebrovascular dysfunction and disease progression [1–6]. CAA-related complications, such as ARIA, have gained attention as a major safety concern associated with newly approved anti-amyloid immunotherapies for early AD [7, 8, 84, 85]. Although ARIA and its subtypes [ARIA-E (edema/effusion) and ARIA-H (microhemorhage/cortical superficial siderosis)] were originally defined in patients undergoing immunotherapy, spontaneous ARIA has also been reported, albeit rarely, in placebo participants and in the context of CAA-related inflammation (CAA-ri) [7–9, 15, 84, 86–92]. This provides a clinical parallel to the spontaneous ARIA events observed in geriatric SQMs with advanced CAA, as reported previously and further characterized in the current study. These observations support the use of this NHP model as a translational platform for investigating pathogenesis and longitudinal trajectories of ARIA, as well as imaging biomarkers relevant to both spontaneous and treatment-associated ARIA in humans [26]. SQMs also offer the opportunity to evaluate iron-related susceptibility changes associated with CAA that may accompany or precede ARIA. The present *in vivo* study employed quantitative *R*_2_*, sensitive to iron-related magnetic field inhomogeneities, to monitor age-related and region-specific brain pathology in SQMs with varying CAA severity, in parallel with characterization of ARIA.

Elevated *R_2_ and R_2_\** have been shown to reflect higher regional iron concentrations and to be associated with cognitive decline related to both aging and neurodegenerative diseases [41, 42, 46, 53, 61, 62, 93]. In line with human findings, whole-brain *R*_2_* increased with age across the SQM lifespan [94]. Total GM *R*_2_* demonstrated a positive cross-sectional association with age, likely reflecting the combined effects of elevated iron burden and cerebrovascular dysfunction driven by CAA in geriatric SQMs. As Aβ deposits accumulate in leptomeningeal and cortical vessels, they contribute to vascular stiffening and lumen narrowing [3, 95], leading to dysregulated blood flow, regional hypoperfusion, and increased deoxyhemoglobin. These changes manifest as hypointense, vessel-like signal patterns on *T*_2_*-w GE images and elevated GM *R*_2_* [51–53]. Consistent with this mechanism, we observed hypointensities along vascular trajectories in our geriatric SQM cohorts, possibly indicative of localized hypoxia and hypoperfusion [51, 54, 96]. These vascular impairments may further exacerbate downstream iron dysregulation and augment magnetic susceptibility [31, 49, 54].

Moreover, iron can accumulate within Aβ deposits, which bind and retain metal ions within their insoluble cores, increasing local magnetic susceptibility [97]. The vascular dysfunction and hypoxia described above further elevate *R*_2_* through an additive hypoxia-driven BOLD effect layered on top of the iron-related signal [53]. Notably, CAA-related structural weakening of vessel walls predisposes affected vessels to microhemorrhages and hemosiderin deposition, producing strong paramagnetic effects [42]. Neuroglial responses, including microglial ferritin upregulation and iron sequestration in perivascular and plaque-adjacent areas, may further amplify susceptibility [98–101].Together, these iron-related and hypoxia-driven mechanisms likely shaped the observed anatomical region- and age-dependent GM *R*_2_* increases in SQMs.

To identify the neuropathological features of age-related *R*_2_* changes, we conducted extensive MRI-histopathology assessment in the late geriatric cohort. Vessel-like hypointensities on *T*_2_*-w MGE scans colocalized with larger CAA-positive vessels characterized by amyloid-iron accumulation, perivascular microgliosis (including iron-sequestering microglia occasionally seen adjacent to vascular Aβ deposits), and reactive astrocytes, along with BBB disruption evidenced by extravasated plasma proteins, together providing histological support for the mechanisms described above. A possible alternative explanation for the hypointense vessels is a physiological imaging artifact rather than true pathology; however, this is less likely, because the animals were continuously monitored by a veterinarian throughout each scan, with vital signs including oxygen saturation, respiration, and body temperature maintained within an appropriate range. As an additional indicator of vascular fragility in this model, we examined ARIA-H. Punctate *T*_2_*-w hypointensities consistent with ARIA-H were observed in several geriatric SQMs and confirmed histologically by hemosiderin within CAA-positive vessels and surrounding parenchyma. Microhemorrhages were commonly accompanied by clusters of microglia and astrocytes. ARIA-H predominantly involved caudal brain regions. Cortical superficial siderosis was also documented in one subject, further underscoring the utility of this model.

Building upon whole-brain observations, we conducted sub-anatomical analysis to delineate and characterize regional *R*_2_* trajectories. The high-resolution 3D isotropic MGE volumes facilitated accurate two-stage (linear followed by nonlinear) intersubject registration and the delineation of GM and WM subregions using an in-house-developed atlas . Regional profiling revealed four distinct *R*_2_* trajectories: three monotonic increase patterns (continuous, early-onset, and late-onset) and a fourth characterized by reversal in later life. Continuous increases predominated in parietal and cingulate cortices, early-onset increases in occipital and temporal cortices, and late-onset changes in prefrontal and frontal cortices.

Several aspects of the known spatial distribution of Aβ and CAA provide context for these profiles. *R*_2_* elevations have been reported in temporal and occipital lobes in AD patients compared to healthy controls [41]. In humans, CAA tends to involve posterior lobar regions, particularly the occipital lobes [102]. In SQMs, caudal brain regions, predominantly the occipital cortex, appear to contain mainly fibrillar vascular Aβ deposits, consistent with our observations [25, 30]. Although total amyloid burden has been reported to be higher in rostral regions such as the prefrontal and frontal cortex, these areas harbor both CAA and parenchymal plaque-like Aβ deposits, with only a subset of plaques being congophilic [25, 30, 103]. This pattern suggests that the posterior fibrillar vascular pathology, rather than non-congophilic parenchymal plaques, may have contributed to the observed regional *R*_2_* trajectories. Further studies are needed to clarify the onset and topography of Aβ-related pathology in this NHP model.

Deep GM structures, such as the globus pallidus, exhibited early-onset *R*_2_* increases, consistent with their high physiological iron content [104]. A trend toward increased *R*_2_* in the caudate nucleus and a non-significant, variable pattern in the thalamus were also observed, whereas the putamen exhibited an initial decline from the young to early geriatric groups followed by a late-life increase that was not significant cross-sectionally but was significant longitudinally. These patterns mirror human lifespan *R*_2_* trajectories [40] and align with reports of elevated basal ganglia *R*_2_* in AD [41] and *R*_2_ in CAA [42]. Hence, our findings extend these observations to a spontaneous NHP model of vascular pathology, underscoring the utility of quantitative *R*_2_* as a sensitive marker to monitor basal ganglia iron storage and age-related accumulation [105].

Interestingly, the hippocampus showed a reversal in *R*_2_*. Clinical studies have reported lower hippocampal *R*_2_ in AD despite increased ferritin iron [106], an effect attributed to tissue loss and microstructural changes. Because pronounced hippocampal atrophy is rarely reported in NHPs, the observed reversal may reflect species-specific biology, differential microstructural alterations, or limited statistical power. This finding motivates targeted longitudinal studies integrating volumetry, QSM, *R*_2_*, and histopathology to clarify hippocampal susceptibility changes and to bridge NHP-human differences.

Building on GM findings, we also observed an age-associated reversal in total WM *R*_2_*, modeled with a quadratic function. Unlike GM, this non-linear trajectory in WM likely reflects progressive myelin breakdown and fluid influx during late aging, which override earlier maturation effects, including ongoing myelination and modest iron accumulation [63]. Disruption of compact, iron- and lipid-rich myelin and oligodendrocyte-associated iron alterations increase tissue water, prolong *T*_2_*, and lower *R*_2_*, consistent with demyelination and axonal degeneration [60]. Thus, myelin loss, altered iron handling, and edema in WM likely contributed to the *R*_2_* decline in late-geriatric SQMs [61]. These observations align with human studies reporting reversal in WM *R*_2_ in AD [46, 64] and decreased WM *R*_2_* in Parkinson’s disease with demyelination, underscoring the impact of WM microstructure on *R*_2_* metrics [62]. Atlas-based WM analysis identified *R*_2_* reversal in several tracts, including corona radiata, forceps major, forceps minor, inferior fronto-occipital fasciculus, and longitudinal fasciculus, suggesting selective vulnerability to edema and myelin loss [64]. Since these tracts did not consistently show corresponding hyperintensities on conventional *T*_2_-w/FLAIR imaging, *R*_2_* reversal may itself serve as a complementary and potentially more sensitive macroscopic biomarker, capable of identifying regions undergoing edema-related change that might otherwise remain undetected. By contrast, the anterior commissure and corpus callosum did not exhibit *R*_2_* reversal. Unlike the tracts with reversal, no clear edema-like lesions were detected in these regions in geriatric SQMs, which may explain the absence of *R*_2_* decline. Further investigation with MRI modalities more sensitive to WM microstructure, such as diffusion MRI, is warranted to characterize these findings. Importantly, several regional GM and WM *R*_2_* alterations identified cross-sectionally were recapitulated in the longitudinal follow-up, despite the limited sample size, further supporting an association between aging and regional *R*_2_* changes in SQMs.

Moreover, we confirmed the propensity of SQMs to develop subcortical spontaneous ARIA-E lesions in late-geriatric animals with severe CAA. These edematous areas were predominantly multifocal, with asymmetric MRI signal hyperintensities involving WM, and to a lesser extent GM, in the parieto-occipital lobes [26], with a prevalence of 58% (7/12) in the late-geriatric SQM cohort. Histopathological assessment of MRI-defined ARIA-E regions revealed pronounced microgliosis and microglial dystrophy, plasma extravasation indicative of BBB disruption, and, in a subset, severe myelin loss and axonal abnormalities consistent with advanced lesion stages. However, in adjacent affected GM, fibrinogen deposits and prominent microgliosis were also accompanied by astrocytosis, T-cell infiltration, and perivascular clustering of astrocytes and microglia around CAA-positive leptomeningeal and parenchymal vessels, suggestive of CAA-associated perivascular inflammation. Furthermore, perivascular inflammation may be particularly implicated in the sequence of events leading to spontaneous microhemorrhages, while the observed axonal disruption raises the possibility that ARIA-E events may carry long-term functional consequences in vulnerable regions. These histopathological findings provide further insights into tissue changes that may underlie treatment-associated ARIA-E in patients [7, 107].

Spontaneous ARIA appears more prominent in SQMs than in humans, warranting future studies to investigate how pre-existing CAA and associated inflammation may influence the response to anti-amyloid therapies. Interestingly, a subset of spontaneous ARIA events in this model may resemble CAA-ri, with overlapping radiological and pathological features including lymphocytic infiltrates, perivascular macrophages, activated microglia and astrocytes, and evidence of BBB disruption [14, 15, 92, 107–110]. However, these events appear to have predominantly asymptomatic clinical manifestations compared with CAA-ri, a clinically overt inflammatory form of CAA [108–110].

Several limitations should be acknowledged. Histological assessment was restricted to the late geriatric cohort because of the scarcity of young and early geriatric SQM brain tissue. Longitudinal *T*_2_*-w MGE and *T*_2_-w imaging, despite the limited number of animals, partly mitigated this limitation by enabling repeated assessment of vascular and hemorrhagic changes over time without reliance on limited histological specimens. Although no overt behavioral deficits were observed, cognitive and sensory performance was not systematically assessed.

## Conclusion

This study provides the first *in vivo* evidence linking CAA-related pathology and *R*_2_* alterations, alongside the characterization of spontaneous ARIA. Here, we document that aging squirrel monkeys develop spontaneous CAA accompanied by iron-associated microvascular pathology, ARIA-E, and ARIA-H sharing key imaging and histopathological features with those reported in humans [8, 10–12, 25, 26, 92]. High-resolution MGE served a dual purpose in this model, providing anatomical guidance and unbiased quantitative *R*_2_* mapping sensitive to both iron and edema, while revealing age-dependent, region-specific *R*_2_* trajectories in GM and WM. Although regional brain iron accumulation is known to increase with age [31], the divergent trajectories observed across regions are informative in their own right, suggesting differential regional vulnerability over the course of aging. Several iron-related pathology substrates likely contributed to these trajectories, including hemosiderin originating from microhemorrhages and superficial siderosis, ferritin-bound storage iron, perivascular inflammatory responses and amyloid -associated iron [31]. Notably, *R*_2_* also showed late-life reversal in tracts prone to edema and myelin loss. This reversal may reflect an unanticipated sensitivity of *R*_2_* to excess tissue fluid associated with edema, capturing a transient signal that conventional *T*_2_-w/FLAIR imaging can miss in otherwise normal-appearing regions, particularly as WM integrity declines with advancing pathology. Furthermore, SQMs presented a valuable opportunity to evaluate ARIA manifestation in association with the brain pathology correlates, an area that remains understudied, particularly in the case of transitory ARIA-E. Another strength of the study was the contemporaneous assessment of tissue integrity within MRI-identified affected regions. MRI-histopathology correlations demonstrated that *T*_2_*-w hypointensities corresponded to CAA-positive, iron-laden vessels with perivascular gliosis and BBB disruption, and, in some cases, evidence of ARIA-H microhemorrhages, consistent with age-associated cerebrovascular dysfunction. ARIA-E hyperintensities were marked by reactive gliosis, plasma extravasation, and, in more advanced lesions, myelin and axonal injury.

Building on the findings presented here, future longitudinal studies with larger SQM cohorts integrating *in vivo* neuroimaging with biofluid biomarkers, cognitive measures, and neuropathological signatures will help address gaps in understanding the long-term trajectories and consequences of CAA and ARIA and further advance the development of translational biomarkers and therapeutic targets. Overall, the current research supports the use of quantitative *R*_2_* and conventional *T*_2_-w/FLAIR imaging as complementary candidate biomarkers for CAA detection and characterization of CAA-related injury, with the goal of enhancing antemortem diagnostic capacity, improving ARIA risk stratification, while further informing treatment strategies for patients with AD and CAA.

## Funding

This research was supported by the following grants from the National Institute of Health: R01NS128190 (H.S.), R21NS153418 (H.S.), U24NS135568 (Y.W., T.W.), R33AG073305 (H.S., W.D.H), P40OD010938 Squirrel Monkey Breeding and Research Resource (W.D.H). The MD Anderson Cancer Center Small Animal Imaging Facility is partially supported by National Cancer Institute Grant P30CA016672.

## Acknowledgements

We sincerely thank Dr. Sarah Kezar, Elizabeth Lindemann, and the clinical and husbandry staff at the Keeling Center for Comparative Medicine and Research at UT MD Anderson Cancer Center for maintaining the SQM colony. We thank Jorge De La Cerda and Jennifer Myer from the Small Animal Imaging Facility at UT MD Anderson Cancer Center for their technical assistance with small animal imaging and data acquisition. We extend our appreciation to Krystal Garrow, Erica Moore, and the clinical and husbandry staff from the Department of Veterinary Medicine and Surgery at UT MD Anderson Cancer Center for their assistance with animal care and husbandry. We acknowledge the contributions of Dr. Henry Rusinek and Artem Mikheev for the design and development of FireVoxel. We thank Thomas Genovese for his help with initial MRI analysis. We gratefully appreciate Mark Alu from the NYU Langone’s Experimental Pathology Research Laboratory for his technical assistance with brain tissue processing.

## Declaration of Interest

The authors declare no competing financial interests.

---

**Supplementary Figure S1.**
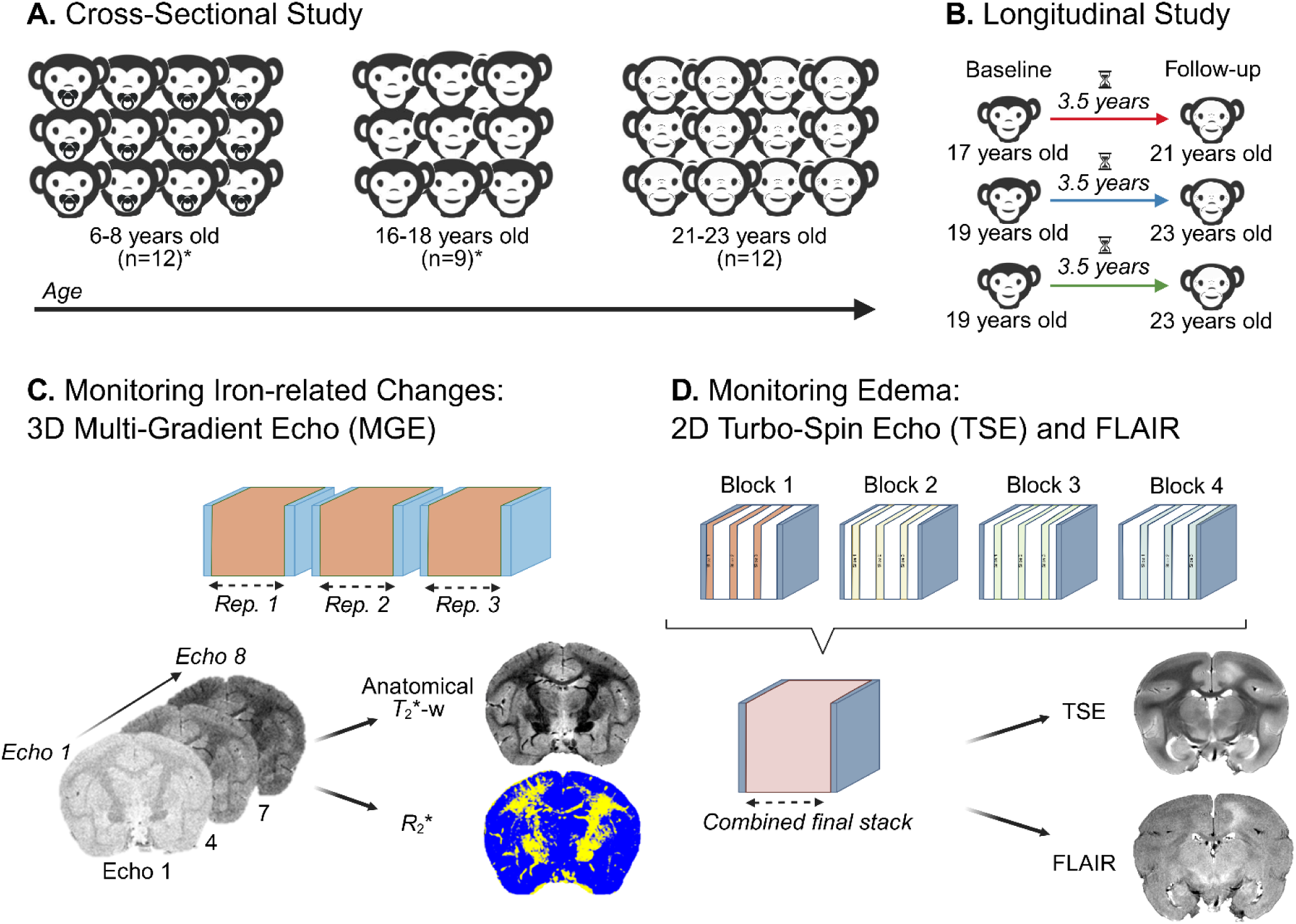
Study design and MRI acquisitions. **(A)** The study employed a cross-sectional design with three SQM age cohorts, number of subjects (33): young (7.10 ± 0.70 years, n = 12), early geriatric (17.61 ± 0.65, n = 9), and late geriatric (21.95 ± 0.96 years, n = 12). **(B)** A longitudinal design monitored three individual subjects through the geriatric period (approximately 17-23 years old) with MRI sessions separated by 3.5 years. **(C)** Workflow for 3D MGE *T*_2_*-w imaging at 200 µm isotropic resolution. A maximum of three repetitions (NR ≤ 3) were acquired if animals remained physiologically stable under anesthesia. Eight echoes were utilized to generate echo-averaged *T*_2_*-w anatomical images with enhanced SNR and contrast-to-noise ratios (CNR) via MRtrix3, and voxel-wise quantitative *R*_2_* map (*R*_2_* = 1/*T*_2_*) derived by mono-exponential fitting in FireVoxel. Both datasets were aligned and transformed to VALiDATe29 template using ITK-SNAP and ANTs for unbiased analysis of iron-related changes across aging groups. **(D)** A 2D multi-slice acquisition strategy designed to enhance detection of edema and fluid-related changes during aging. To ensure high-resolution whole-brain coverage, data were acquired in four interleaved blocks (800 µm slices, 200 µm in-plane, 2.4 mm slice spacing) and subsequently merged into a single contiguous volume (MRtrix3). *T*_2_-w TSE and FLAIR sequences utilized identical spatial parameters, with FLAIR incorporating an inversion pulse (TI = 1557 ms) to suppress CSF. An asterisk (*) for the number of subjects in each cohort indicates that both 3D *T*_2_*-w MGE and 2D *T*_2_-w sequences were acquired; for full breakdown of study population and imaging protocol see **Supplementary Table 1 ST1.**

**Supplementary Figure S2:**
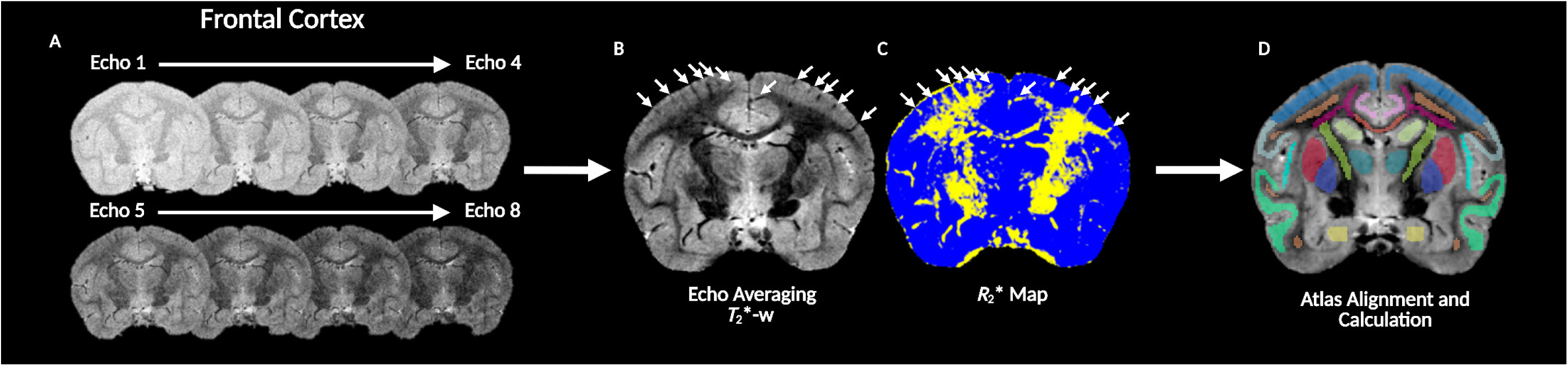
Dual-purpose MGE acquisition: anatomical guidance and quantitative *R*_2_* mapping. **(A)** Echo-dependent SNR decline consistent with *T*_2_* decay, illustrating the complementary trade-off between SNR and GM/WM contrast across echoes. **(B)** Multi-echo-averaged volume showing markedly improved SNR, CNR, and reduced signal loss near air/tissue interfaces. **(C)** Pixel-wise *R*_2_* map providing an absolute, acquisition-independent iron-sensitive biomarker enabling direct cross-cohort comparisons. **(D)** Superior tissue contrast enabling fine anatomical delineation and atlas refinement

**Supplementary Figure S3:**
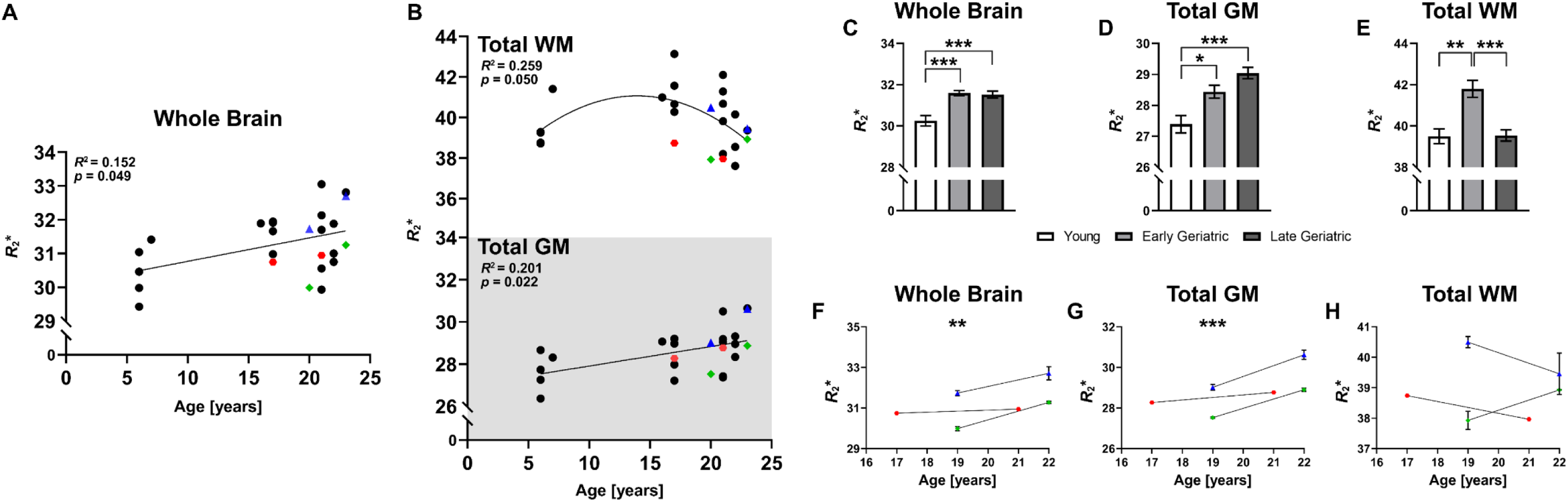
Age-related whole-brain and tissue-specific *R*_2_* trajectories. **(A)** Whole-brain *R*_2_* increases significantly with age. **(B)** GM *R*_2_* shows a significant positive age correlation, while WM *R*_2_* follows a curvilinear trajectory, with an initial increase followed by reversal at later ages. Three longitudinally monitored subjects (color-coded paired symbols) are overlaid across panels, corroborating cross-sectional trends. Statistical Analysis: Pearson’s Correlation Coefficient, Linear and Quadratic Regressions. \**p* < 0.05; \*\**p* < 0.01; \*\*\**p* < 0.001. **(C–E)** Group-wise cross-sectional comparisons of whole-brain **(C)**, total GM **(D)**, and total WM **(E)** *R*_2_* across young (n = 5), early geriatric (n = 6), and late geriatric (n = 12) cohorts confirm the trends observed in panels A and B. Statistical Analysis: One-way ANOVA with Bonferroni correction; see full statistics in **Supplementary Table ST3**. Significance: \**p* < 0.05; \*\**p* < 0.01; \*\*\**p* < 0.001. **(F–H)** Longitudinal trajectories of whole-brain **(F)**, total GM **(G)**, and total WM **(H)** *R*_2_* in three individually monitored SQMs (color-coded), illustrating within-subject changes over time that corroborate the cross-sectional patterns and show progressive increases in whole-brain and GM *R*_2_*, with late-life reversal in WM consistent with the quadratic fit in panel B. Statistical Analysis: Paired *t*-test throughout; see full statistics in **Supplementary Table ST4**. Significance: \**p* ≤ 0.05; \*\**p* ≤ 0.01; \*\*\**p* ≤ 0.001.

**Supplementary Figure S4:**
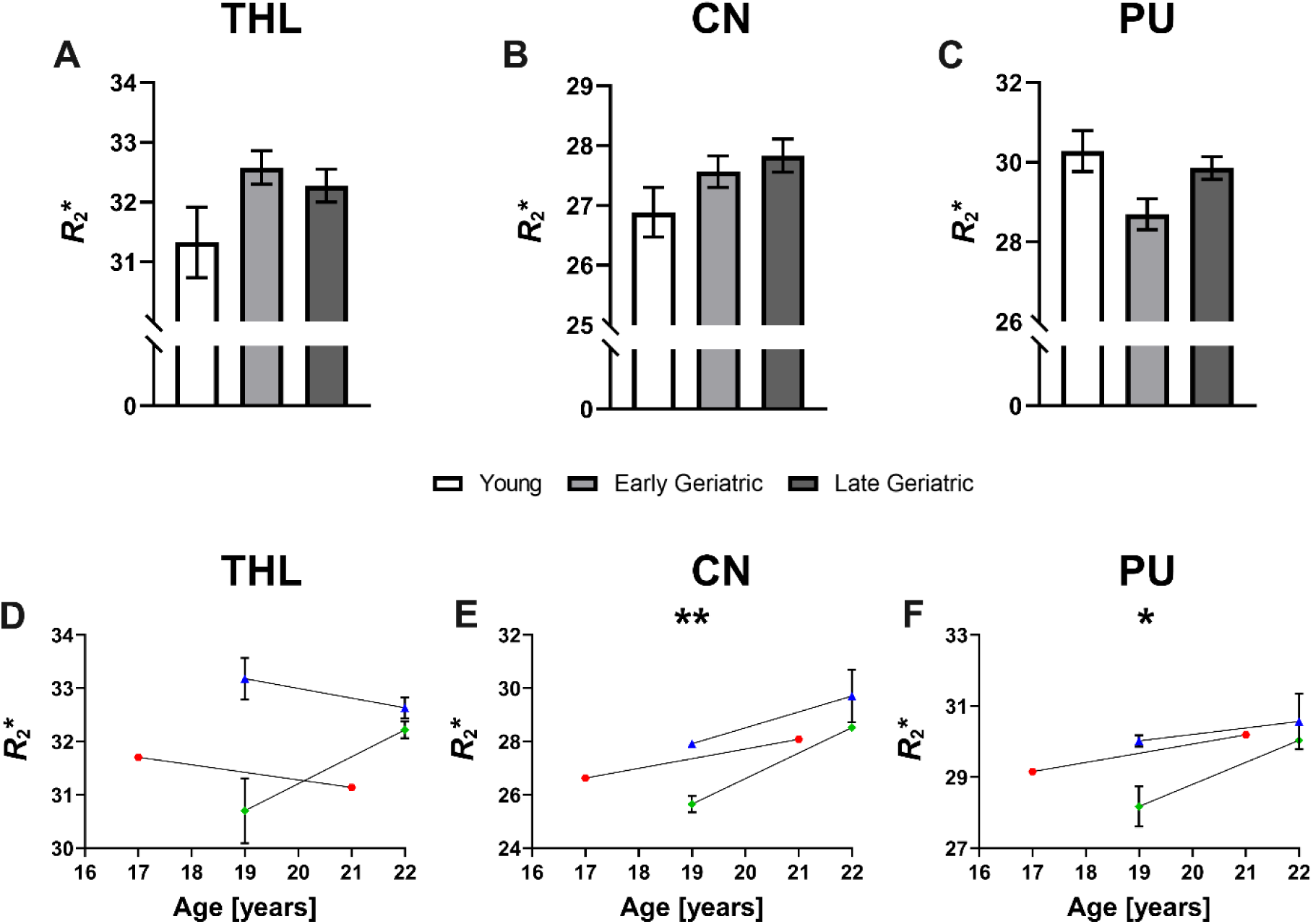
Age-related *R*_2_* changes across deep gray matter structures. (A–C) Group-wise cross-sectional *R*_2_* values (mean ± SEM) in Thalamus (THL), Caudate Nucleus (CN), and Putamen (PU) across young, early geriatric, and late geriatric SQM cohorts. THL shows a non-significant variable pattern across age groups, CN exhibits a progressive age-related increase, and PU displays an initial decline from young to early geriatric followed by a non-significant late-life increase. Statistical Analysis: One-way ANOVA with Bonferroni correction; see full statistics in **Supplementary Table ST3**. Significance: \**p* ≤ 0.05; \*\**p* ≤ 0.01; \*\*\**p* ≤ 0.001. **(D–F)** Longitudinal *R*_2_* trajectories in THL **(D)**, CN **(E)**, and PU **(F)** for three individually monitored SQMs, based on two geriatric time points (early and late), illustrating within-subject changes that qualitatively complement the cross-sectional patterns. CN shows a significant upward pattern; THL changes are more heterogenous, and PU shows an increase at the later time point without reaching statistical significance, but directionally consistent with the cross-sectional results. Statistical Analysis: Paired *t*-test throughout; see full statistics for GM and WM regions in **Supplementary Table ST4.** Significance: \**p* ≤ 0.05; \*\**p* ≤ 0.01; \*\*\**p* ≤ 0.001.

**Supplementary Figure S5.**
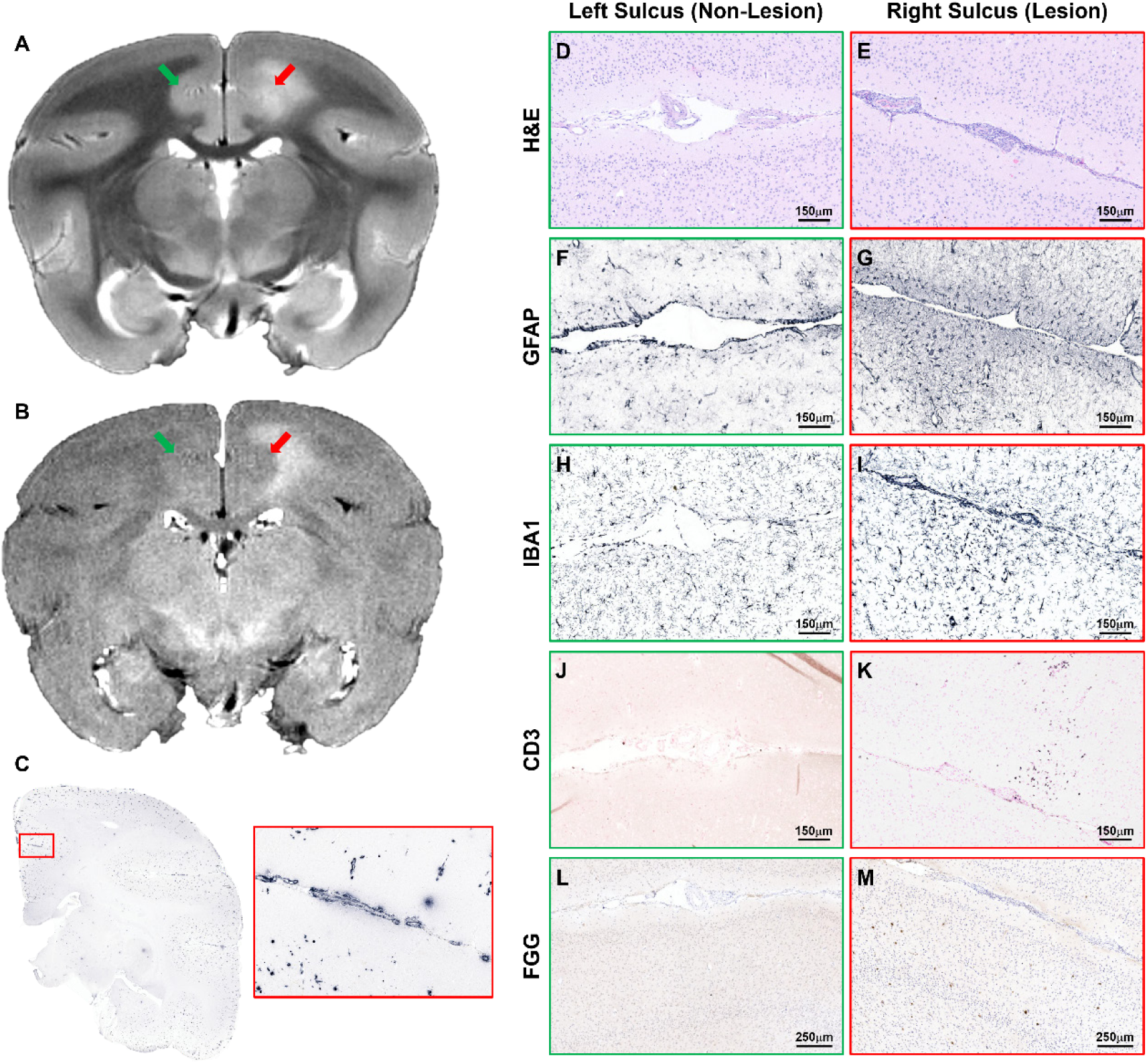
Histopathological characterization of ARIA-E signal abnormality in the cingulate sulcus and adjacent GM. **(A)** *In vivo T*_2_-w MRI revealed a hyperintense lesion (same subject and lesion as **in** Figure 7) extending into the cingulate sulcus and adjacent GM within the fronto-parietal lobe of the right hemisphere (red arrow); contralateral normal-appearing sulcus is indicated in the left hemisphere (green arrow). **(B)** Corresponding coronal *T*_2_-w FLAIR sequence confirmed the extent of the lesion. **(C)** Low-magnification view of 6E10/4G8 immunostaining, with a higher-magnification inset of the sulcal region, confirming vascular amyloid-β deposition. A representative set of histology images is shown for left hemisphere with normal-appearing GM and sulcus (non-lesion, green boxes) and contralateral right hemisphere with affected GM and sulcus (lesion, red boxes), stained for **(D, E)** H&E, **(F, G)** GFAP, **(H, I)** IBA1, **(J, K)** CD3, and **(L, M)** FGG. In the hyperintense sulcal area, H&E staining showed a cell-rich region associated with leptomeningeal vessels affected by CAA. GFAP and IBA1 immunohistochemistry demonstrated prominent astrocytosis and microgliosis in close proximity to sulcal leptomeningeal vessels, indicative of focal CAA-associated perivascular inflammation. CD3 immunostaining revealed infiltration of T lymphocytes within the parenchyma of the edematous GM area, whereas no CD3-positive cells were detected in the contralateral unaffected hemisphere. Fibrinogen immunoreactivity was present multifocally along microvessels in the ARIA-E affected GM region, consistent with focal BBB disruption. Scale bars are as indicated.

**Supplementary Figure S6.**
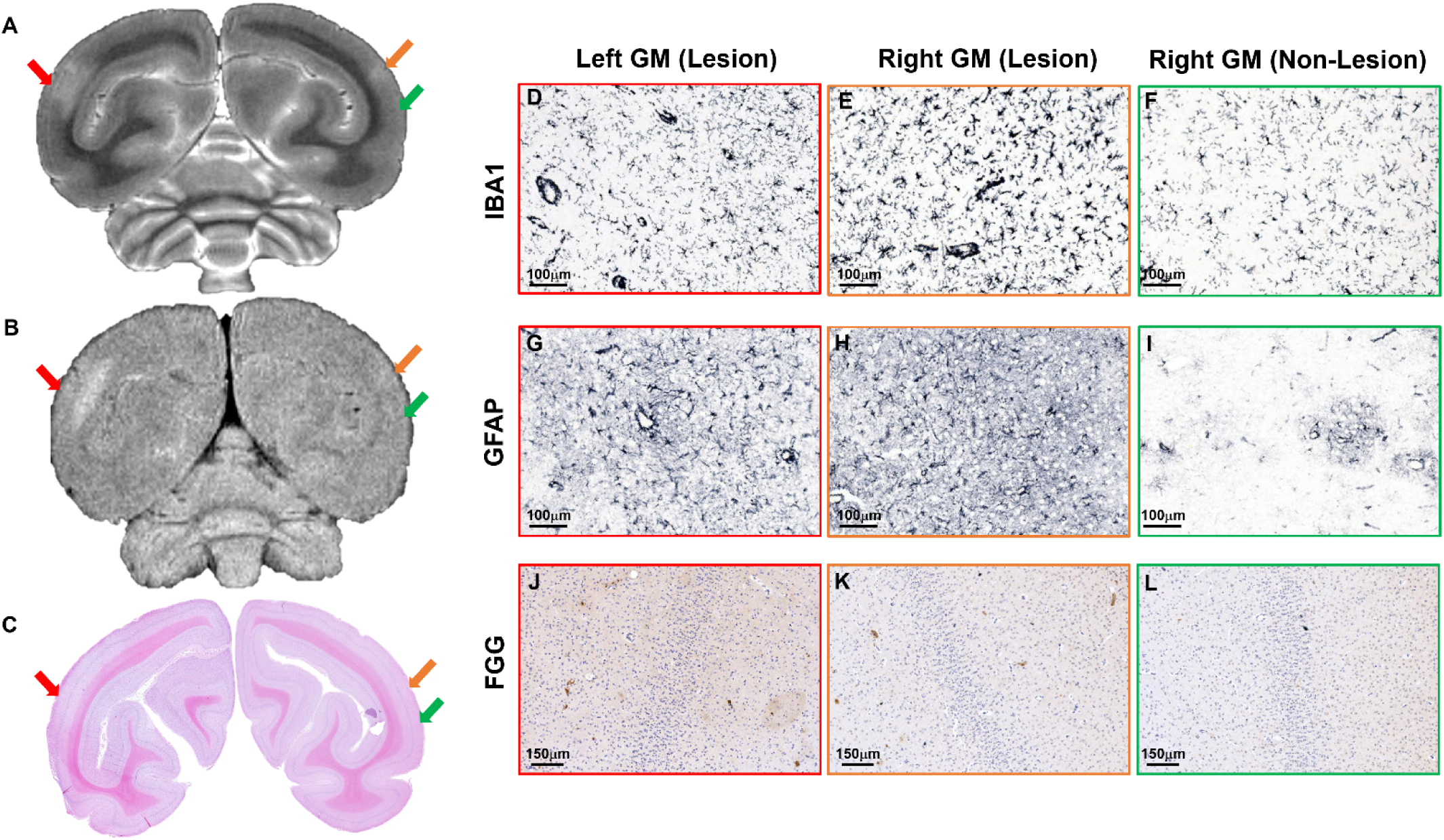
Histopathological characterization of ARIA-E signal abnormality in adjacent occipital GM. **(A)** *In vivo T*_2_-w MRI revealed a hyperintense lesion (same subject and lesion as **in** Figure 8) extending into the adjacent GM within the occipital lobe of the left hemisphere (red arrow), in addition to a smaller hyperintense area detected in the right hemisphere (orange arrow); contralateral normal-appearing GM area is indicated in the right hemisphere (green arrow). **(B)** Corresponding coronal *T*_2_-w FLAIR sequence confirmed the extent of the lesion. **(C)** Matched histological section stained with H&E. A representative set of histology images is shown for both hemispheres with ARIA-E affected GM (lesion, red and orange boxes) and right hemisphere with normal-appearing GM region (non-lesion, green boxes) stained for **(D-F)** IBA1, (**G-I)** GFAP, **(J-L)** FGG. Both hyperintense GM regions demonstrated prominent GFAP astrocytosis and IBA1 microgliosis associated with perivascular clustering of reactive astrocytes and microglia. Multifocal fibrinogen deposits were evident in both ARIA-E GM regions, indicative of focal BBB disruption. Scale bars are as indicated.

**Supplementary Table ST1:** Characteristics of the study population and imaging protocol. This table lists individual SQM subjects with their identifiers (ID), age at each scan, and imaging protocols used: *T*_2_*-w MGE, *T*_2_-w TSE/FLAIR, or both. **Age distribution by cohort:** young: 6.11-8.13 years; mean ± SD = 7.10 ± 0.70; median = 7.05; n = 12; CV = 10%. Early geriatric: 16.54-18.55 years; mean ± SD = 17.50 ±0.65; median = 17.44; n = 9; CV = 4%. Late geriatric: 20.97-23.58 years; mean ± SD = 21.95 ± 0.96; median = 21.83; n = 12; CV = 4%. **Age distribution** is summarized by cohort (young, early geriatric, late geriatric). Values are reported as minimum to maximum, mean ± standard deviation (SD), median, number of subjects (n), and coefficient of variation (CV). An asterisk (*) marks subject ID 14 in both the early and late geriatric cohorts, indicating a longitudinal MRI assessment.

| Cohort | Subject ID | Age | $T_2^*$-w (MGE) | $T_2$-w (TSE/FLAIR) |
| --- | --- | --- | --- | --- |
| Young<br>(n = 12) | 1 | 6.53 | X |  |
|  | 2 | 6.52 | X |  |
|  | 3 | 6.52 | X |  |
|  | 4 | 6.50 | X |  |
|  | 5 | 7.09 | X |  |
|  | 6 | 7.07 |  | X |
|  | 7 | 7.02 |  | X |
|  | 8 | 7.75 |  | X |
|  | 9 | 7.94 |  | X |
|  | 10 | 8.04 |  | X |
|  | 11 | 6.11 |  | X |
|  | 12 | 8.13 |  | X |
| Early Geriatric<br>(n = 9) | 13 | 17.45 | X | X |
|  | 14* | 17.40 | X |  |
|  | 15 | 17.44 | X | X |
|  | 16 | 17.43 | X |  |
|  | 17 | 16.71 | X |  |
|  | 18 | 18.55 | X |  |
|  | 21 | 16.54 |  | X |
|  | 20 | 17.59 |  | X |
|  | 19 | 18.36 |  | X |
| Late Geriatric<br>(n = 12) | 22 | 21.76 | X | X |
|  | 23 | 21.46 | X | X |
|  | 24 | 23.58 | X | X |
|  | 25 | 23.57 | X | X |
|  | 26 | 20.98 | X | X |
|  | 27 | 21.90 | X | X |
|  | 28 | 21.20 | X | X |
|  | 29 | 21.99 | X | X |
|  | 14* | 20.97 | X | X |
|  | 30 | 23.03 | X | X |
|  | 31 | 21.92 | X | X |
|  | 32 | 21.01 | X | X |

**Supplementary Table ST2:** Characteristics of the longitudinal study population and imaging protocol. This table lists the subset of SQM subjects that underwent longitudinal MRI assessment, with their identifiers (ID), age at each scan, and imaging protocols used: *T*_2_*-w MGE, *T*_2_-w TSE/FLAIR, or both. The table illustrates within-subject aging trajectories and changes in imaging coverage across sessions.

| Subject ID | Time point | Age | $T_2^*$-w (MGE) | $T_2$-w (TSE/FLAIR) |
| --- | --- | --- | --- | --- |
| 14 | Baseline | 17.4 | X |  |
|  | Follow-up | 20.97 | X | X |
| 24 | Baseline | 19.98 | X |  |
|  | Follow-up | 23.58 | X | X |
| 25 | Baseline | 19.95 | X |  |
|  | Follow-up | 23.57 | X | X |

**Supplementary Table ST3:**
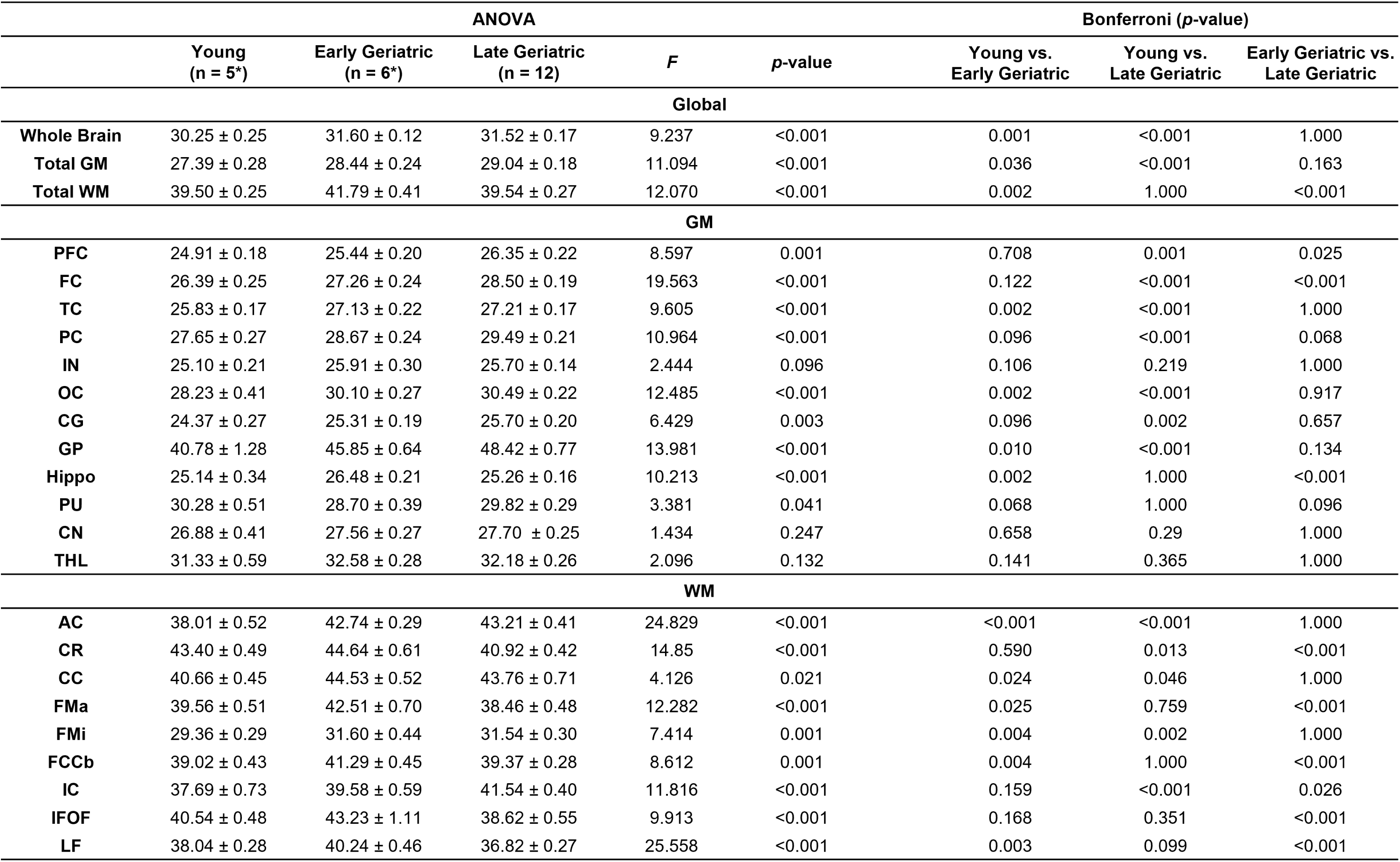
Cross-sectional and head-to-head comparisons of regional *R*_2_*\** values. Cohort-based *R*_2_*\** values are presented as mean ± SEM. n denotes the number of subjects. An asterisk (*) for the number of subjects in each cohort indicates that only 3D *T*_2_*-w MGE sequences were acquired; for full breakdown of study population and imaging protocol see **Supplementary Table 1 ST1. Regions:** Total GM, Total gray matter; Total WM, Total white matter; PFC, Prefrontal Cortex; FC, Frontal Cortex; TC, Temporal Cortex; PC, Parietal Cortex; IN, Insula; OC, Occipital Cortex; CG, Cingulate Cortex; GP, Globus Pallidus; Hippo, Hippocampus; PU, Putamen; CN, Caudate Nucleus; THL, Thalamus; AC, Anterior Commissure; CR, Corona Radiata; CC, Corpus Callosum; FMa, Forceps Major; FMi, Forceps Minor; FCCb, Fiber Extension of Corpus Callosum body; IC, Internal Capsule; IFOF, Inferior Fronto-Occipital Fasciculus; LF, Longitudinal Fasciculus. Statistical Analysis: One-way ANOVA with Bonferroni post-hoc correction. *F* indicates the F-statistic. Significance: \**p* ≤ 0.05; \*\**p* ≤ 0.01; \*\*\**p* ≤ 0.001.

|  | ANOVA |  |  |  |  | Bonferroni (*p*-value) |  |  |
| --- | --- | --- | --- | --- | --- | --- | --- | --- |
|  | Young<br>(*n* = 5*) | Early Geriatric<br>(*n* = 6*) | Late Geriatric<br>(*n* = 12) | *F* | *p*-value | Young vs.<br>Early Geriatric | Young vs.<br>Late Geriatric | Early Geriatric vs.<br>Late Geriatric |
| **Global** |  |  |  |  |  |  |  |  |
| **Whole Brain** | 30.25 ± 0.25 | 31.60 ± 0.12 | 31.52 ± 0.17 | 9.237 | <0.001 | 0.001 | <0.001 | 1.000 |
| **Total GM** | 27.39 ± 0.28 | 28.44 ± 0.24 | 29.04 ± 0.18 | 11.094 | <0.001 | 0.036 | <0.001 | 0.163 |
| **Total WM** | 39.50 ± 0.25 | 41.79 ± 0.41 | 39.54 ± 0.27 | 12.070 | <0.001 | 0.002 | 1.000 | <0.001 |
| **GM** |  |  |  |  |  |  |  |  |
| **PFC** | 24.91 ± 0.18 | 25.44 ± 0.20 | 26.35 ± 0.22 | 8.597 | 0.001 | 0.708 | 0.001 | 0.025 |
| **FC** | 26.39 ± 0.25 | 27.26 ± 0.24 | 28.50 ± 0.19 | 19.563 | <0.001 | 0.122 | <0.001 | <0.001 |
| **TC** | 25.83 ± 0.17 | 27.13 ± 0.22 | 27.21 ± 0.17 | 9.605 | <0.001 | 0.002 | <0.001 | 1.000 |
| **PC** | 27.65 ± 0.27 | 28.67 ± 0.24 | 29.49 ± 0.21 | 10.964 | <0.001 | 0.096 | <0.001 | 0.068 |
| **IN** | 25.10 ± 0.21 | 25.91 ± 0.30 | 25.70 ± 0.14 | 2.444 | 0.096 | 0.106 | 0.219 | 1.000 |
| **OC** | 28.23 ± 0.41 | 30.10 ± 0.27 | 30.49 ± 0.22 | 12.485 | <0.001 | 0.002 | <0.001 | 0.917 |
| **CG** | 24.37 ± 0.27 | 25.31 ± 0.19 | 25.70 ± 0.20 | 6.429 | 0.003 | 0.096 | 0.002 | 0.657 |
| **GP** | 40.78 ± 1.28 | 45.85 ± 0.64 | 48.42 ± 0.77 | 13.981 | <0.001 | 0.010 | <0.001 | 0.134 |
| **Hippo** | 25.14 ± 0.34 | 26.48 ± 0.21 | 25.26 ± 0.16 | 10.213 | <0.001 | 0.002 | 1.000 | <0.001 |
| **PU** | 30.28 ± 0.51 | 28.70 ± 0.39 | 29.82 ± 0.29 | 3.381 | 0.041 | 0.068 | 1.000 | 0.096 |
| **CN** | 26.88 ± 0.41 | 27.56 ± 0.27 | 27.70 ± 0.25 | 1.434 | 0.247 | 0.658 | 0.29 | 1.000 |
| **THL** | 31.33 ± 0.59 | 32.58 ± 0.28 | 32.18 ± 0.26 | 2.096 | 0.132 | 0.141 | 0.365 | 1.000 |
| **WM** |  |  |  |  |  |  |  |  |
| **AC** | 38.01 ± 0.52 | 42.74 ± 0.29 | 43.21 ± 0.41 | 24.829 | <0.001 | <0.001 | <0.001 | 1.000 |
| **CR** | 43.40 ± 0.49 | 44.64 ± 0.61 | 40.92 ± 0.42 | 14.85 | <0.001 | 0.590 | 0.013 | <0.001 |
| **CC** | 40.66 ± 0.45 | 44.53 ± 0.52 | 43.76 ± 0.71 | 4.126 | 0.021 | 0.024 | 0.046 | 1.000 |
| **FMa** | 39.56 ± 0.51 | 42.51 ± 0.70 | 38.46 ± 0.48 | 12.282 | <0.001 | 0.025 | 0.759 | <0.001 |
| **FMi** | 29.36 ± 0.29 | 31.60 ± 0.44 | 31.54 ± 0.30 | 7.414 | 0.001 | 0.004 | 0.002 | 1.000 |
| **FCCb** | 39.02 ± 0.43 | 41.29 ± 0.45 | 39.37 ± 0.28 | 8.612 | 0.001 | 0.004 | 1.000 | <0.001 |
| **IC** | 37.69 ± 0.73 | 39.58 ± 0.59 | 41.54 ± 0.40 | 11.816 | <0.001 | 0.159 | <0.001 | 0.026 |
| **IFOF** | 40.54 ± 0.48 | 43.23 ± 1.11 | 38.62 ± 0.55 | 9.913 | <0.001 | 0.168 | 0.351 | <0.001 |
| **LF** | 38.04 ± 0.28 | 40.24 ± 0.46 | 36.82 ± 0.27 | 25.558 | <0.001 | 0.003 | 0.099 | <0.001 |

**Supplementary Table ST4.** Longitudinal changes in regional *R*_2_* values. *R*_2_* values are presented as mean ± SEM at baseline and follow-up MRI sessions. n denotes the number of subjects. For full breakdown of study population imaging protocol see **Supplementary Table 1 ST1. Regions**: Total GM, Total gray matter; Total WM, Total white matter; PFC, Prefrontal Cortex; FC, Frontal Cortex; TC, Temporal Cortex; PC, Parietal Cortex; IN, Insula; OC, Occipital Cortex; CG, Cingulate Cortex; GP, Globus Pallidus; Hippo, Hippocampus; PU, Putamen; CN, Caudate Nucleus; THL, Thalamus; AC, Anterior Commissure; CR, Corona Radiata; CC, Corpus Callosum; FMa, Forceps Major; FMi, Forceps Minor; FCCb, Fiber Extension of Corpus Callosum body; IC, Internal Capsule; IFOF, Inferior Fronto-Occipital Fasciculus; LF, Longitudinal Fasciculus. Statistical Analysis: paired *t*-test. *t*, T-test statistics. *Z^a^*, *Z* statistics *(Wilcoxon Signed Rank Test). df*, Degrees of freedom. Significance: \**p* ≤ 0.05; \*\**p* ≤ 0.01; \*\*\**p* ≤ 0.001.

|  | Baseline<br>(n = 3) | Follow-up<br>(n = 3) | *t*/*Z* | *p*-value |
| --- | --- | --- | --- | --- |
| **Global** |  |  |  |  |
| **Whole Brain** | 30.84 ± 0.34 | 31.83 ± 0.34 | -5.439 | 0.002 |
| **Total GM** | 28.27 ± 0.29 | 29.61 ± 0.37 | -8.238 | <0.001 |
| **Total WM** | 39.14 ± 0.50 | 39.83 ± 0.34 | 0.242 | 0.816 |
| **GM** |  |  |  |  |
| **PFC** | 25.41 ± 0.28 | 26.70 ± 0.20 | -8.02 | <0.001 |
| **FC** | 27.16 ± 0.21 | 28.64 ± 0.25 | -7.55 | <0.001 |
| **TC** | 26.78 ± 0.17 | 27.74 ± 0.34 | -4.2 | 0.006 |
| **PC** | 28.52 ± 0.36 | 30.04 ± 0.55 | -6.78 | <0.001 |
| **IN** | 25.44 ± 0.17 | 25.57 ± 0.23 | -0.47 | 0.658 |
| **OC** | 29.66 ± 0.38 | 31.26 ± 0.63 | -5.18 | 0.002 |
| **CG** | 24.98 ± 0.25 | 25.74 ± 0.36 | -1.34 | 0.227 |
| **GP** | 48.75 ± 0.83 | 52.86 ± 2.00 | -1.52<sup>a</sup> | 0.128 |
| **Hippo** | 25.61 ± 0.14 | 25.50 ± 0.30 | 0.48 | 0.644 |
| **PU** | 29.26 ± 0.40 | 30.30 ± 0.38 | -1.36<sup>a</sup> | 0.040 |
| **CN** | 26.76 ± 0.43 | 28.98 ± 0.46 | -4.64<sup>a</sup> | 0.004 |
| **THL** | 31.90 ± 0.51 | 32.26 ± 0.25 | -0.82 | 0.444 |
| **WM** |  |  |  |  |
| **AC** | 40.33 ± 0.62 | 43.72 ± 0.79 | -2.03 | 0.043 |
| **CR** | 41.94 ± 0.81 | 39.48 ± 0.52 | 2.72 | 0.035 |
| **CC** | 42.91 ± 0.46 | 42.59 ± 1.25 | -0.18 | 0.864 |
| **FMa** | 38.53 ± 0.51 | 38.71 ± 1.33 | -0.34 | 0.735 |
| **FMi** | 29.12 ± 0.36 | 32.13 ± 0.69 | -4.35 | 0.005 |
| **FCCb** | 38.96 ± 0.59 | 38.52 ± 0.27 | 0.18 | 0.862 |
| **IC** | 39.09 ± 0.98 | 42.59 ± 0.43 | -4.57 | 0.004 |
| **IFOF** | 38.54 ± 0.30 | 37.83 ± 0.74 | 0.736 | 0.490 |
| **LF** | 37.19 ± 0.43 | 36.07 ± 0.21 | 3.69 | 0.010 |

**Supplementary Table ST5.** Cumulative incidence of ARIA-E in study cohorts. This table outlines the incidence of ARIA-E in the subset of SQMs that underwent *T*_2_-w TSE/FLAIR to monitor the presence of MRI abnormalities; n denotes the number of subjects. An asterisk (*) for the number of subjects in the young and early geriatric cohorts indicates that only 2D *T*_2_-w TSE/FLAIR sequences were acquired; for full breakdown of study population and imaging protocol see **Supplementary Table ST1.**

| Cohort | Age range (years) | Subjects with ARIA-E (n) | Subjects examined (n) | ARIA-E incidence |
| --- | --- | --- | --- | --- |
| Young | 6-8 | 0 | 7* | 0% |
| Early Geriatric | 16-18 | 0 | 5* | 0% |
| Late Geriatric | 21-23 | 7 | 12 | 58.33% |

## Notes

### Competing Interest Statement

The authors have declared no competing interest.

